# Identification and small molecule rescue of mitochondrial dysfunction phenotype which converges across Leigh Syndrome and Huntington’s Disease patient fibroblasts

**DOI:** 10.64898/2026.06.08.730157

**Authors:** Naomi Hartopp, Laura Ellis, Rachel Hughes, Emily Mossman, Ella Simmonite, Anastasia Thoma, Fatima-ezzahra Erichidi, Gauri Bhosale, Alessandro Pristerà, Scott P. Allen, Laura Ferraiuolo, Pamela J. Shaw, Oliver Bandmann, Heather Mortiboys

**Affiliations:** Sheffield Institute for Translational Neuroscience, School of Medicine and Population Health, University of Sheffield, 385a Glossop Road, Sheffield, S10 2HQ, United Kingdom; Astellas Pharma Inc., Frontier Biology; NIHR Sheffield Biomedical Research Centre, Sheffield Teaching Hospitals NHS Foundation Trust, Royal Hallamshire Hospital, Glossop Road, Sheffield S10 2JF, United Kingdom

**Keywords:** Mitochondria, mitochondrial disease, neurodegenerative disease, high content imaging, patient-derived fibroblasts, drug screening

## Abstract

Mitochondrial dysfunction is implicated in a variety of complex neurological disorders. Primary mitochondrial diseases are caused directly by mutations in genes encoding mitochondrial proteins, leading to mitochondrial dysfunction and disease. Mitochondrial dysfunction is also a key contributor to pathogenesis in multiple neurodegenerative diseases. Rescue of mitochondrial function is therefore an attractive therapeutic target in both groups of diseases. In this study, we used primary fibroblasts derived from patients with the primary mitochondrial disease Leigh syndrome (LS) and the neurodegenerative disease Huntington’s disease (HD) to investigate mitochondrial phenotypes in these patients. We used these to identify modifiable measures of mitochondrial phenotype using a high content imaging screen. Despite having distinct underlying disease causes, different mitochondrial phenotypes in LS and HD patient derived cells converged on an imbalance between functional and dysfunctional mitochondria. Through multi-parameter screening of the mitochondrial phenotype we identified the AMPK activator A769662 as a small molecule able to rescue this imbalance in both LS and HD patient derived fibroblasts via different pathways. Our findings indicate that high throughput screening for mitochondrial phenotypes could identify novel therapeutic agents to rescue mitochondrial dysfunction in complex neurological disorders.

**Research in Context:** *Evidence before this study:* Mitochondrial dysfunction is the primary driver of mitochondrial disease and key contributing factor to neurodegenerative disease pathogenesis. Rescuing mitochondrial function is a promising therapeutic strategy, yet strategies to identify mitochondrial modulators in patient cells are limited.

*Added value of this study:* Our in-depth mitochondrial characterisation of Leigh syndrome and Huntington’s disease patient fibroblasts shows the potential for using this methodology to identify mitochondrial therapeutics. We identify a mitochondrial phenotype common across diseases and an AMPK activator capable of rescuing this phenotype.

*Implications of all the available evidence:* Our work extends our understanding of the mitochondrial dysfunction associated with Leigh syndrome and Huntington’s disease and expands the tool set available for identifying modulators of mitochondrial health as potential therapeutics for complex neurological disorders.

## Introduction

Mitochondrial function is fundamental to cellular health and mitochondrial dysfunction is implicated in a variety of complex diseases. Genetic defects affecting the mitochondrial respiratory chain cause a group of rare conditions known as primary mitochondrial diseases which lead to severe neurological symptoms early in life and often shorten life expectancy^1^. Mitochondrial dysfunction is also a contributing mechanism to multiple neurodegenerative diseases which cause a variety of motor and cognitive symptoms usually later in life and may be genetically determined or occur sporadically^2,3^. Rescue of mitochondrial function is therefore a promising option for therapeutic intervention in both mitochondrial and neurodegenerative diseases, yet both groups of disorders still represent high unmet clinical need due to challenges in disease modelling and drug discovery for such complex diseases.

Leigh syndrome (LS) is a primary mitochondrial disease which causes progressive neurological deterioration, often presenting before 2 years of age and causing death before 10 years of age^4^. Symptoms are severe and varied, including developmental delay, respiratory dysfunction, seizures, motor weakness and non-neurological symptoms such as gastrointestinal features including diarrhoea and vomiting. There are currently no approved treatments for LS, although small trials are underway across mitochondrial disease patients, for example with Coenzyme Q10 and Vatiquinone (reviewed in Margo et al 2025^5^). LS is genetically and clinically heterogeneous, caused by mutations in nuclear or mitochondrial DNA, all leading to defects in different complexes of the mitochondrial respiratory chain^5^. More than 100 mutations are reported in *SURF1* causing LS with associated cytochrome C oxidase (COX) deficiency^6–8^. Mice with a homozygous SURF1 knockout recapitulate deficiency in COX activity and activate mitochondrial stress responses but do not develop the same respiratory damage or clinical features of the human disease. In fact, these mice have an increased life span compared to controls^9,10^, highlighting species differences and the need for disease modelling in human tissues for drug discovery. Deficits in respiratory complexes were first reported in LS using measurements from patients’ fibroblasts^11,12^. Fibroblasts from LS patients with mutations in *SURF1* display reduced activity of mitochondrial electron transport chain complexes, reduced respiratory activity and damaged mitochondrial membrane potential^13^. These data from peripheral fibroblasts are recapitulated in an induced pluripotent stem cell (iPSC)-derived neuron and organoid model of LS with *SURF1* mutations, which display damaged mitochondrial respiration^14^. Since patient derived fibroblasts are a proliferative, readily accessible tissue, displaying disease relevant phenotypes consistent with neuronal models of disease, they represent an appealing option for high throughput drug discovery.

Huntington’s disease (HD) is a neurodegenerative disease causing cognitive and motor symptoms such as choreic movements, dysphagia and dementia as well as increasingly recognised peripheral symptoms. HD is monogenic, caused by an autosomal dominant CAG repeat expansion in a single gene encoding the huntingtin (HTT) protein, leading to the loss of striatal medium spiny neurons, and neurodegeneration^15,16^. Although longer repeat expansions can result in juvenile forms of HD, most patients manifest symptoms in their 30s-40s. Despite the singular genetic cause, HD manifests as a complex heterogeneous neurodegenerative disease where multiple mechanisms contribute to disease progression. There is no cure for HD and current treatment options are only symptomatic. Although progress has been made in gene therapy approaches aiming to slow HD progression by lowering the level of mutant huntingtin, this strategy has not been successful. The definitive phase 3 trial for the antisense oligonucleotide Tominerson had to be halted due to safety concerns and lack of efficacy ^17^ and trials of the orally available HTT messenger RNA splicing modulator Branaplam were halted due to unacceptable side effects in a phase 2b study^18^. The recent UniQure gene therapy AMT-130 reportedly improves symptoms, but this has not yet been accepted by the FDA. Alternative approaches aiming to correct downstream effects of mutant huntingtin therefore remain justified and should perhaps be considered again with higher priority than before, given the recent failures of experimental therapies aiming to safely lower mutant huntingtin.

The N-terminal of the HTT protein contains a mitochondrial targeting sequence and interacts with multiple mitochondrial proteins. Mutant HTT alters fission-fusion dynamics, increases susceptibility to the mitochondrial permeability transition pore, impairs mitochondrial protein import, mitochondrial transport and mitophagy^19–24^. Importantly, in post mortem HD brains, mitochondria are lost in the spiny caudate neurons^25^ which selectively degenerate during the disease, and increased oxidative stress is reported in HD patient brains before symptom onset^27^. Many HD studies utilise stem cell derived neuronal models, post mortem tissue from patients and/or long CAG repeat expansion knock-in mice models to study the effect of mutant HTT. These studies have reported altered fission-fusion balance, defective bioenergetics, increased reactive oxygen species production and damage to mitophagy (Reviewed in Sawant et al 2021^24^). Mitochondrial swelling and damage to cristae structure and altered mitochondrial protein levels are reported in HD patient fibroblasts^28^ and fibroblasts from patient with earlier onset of disease display lower respiratory capacity and ATP levels^26^, indicating the presence of disease relevant phenotypes in peripheral cells.

Drug discovery for complex neurological disorders faces significant challenges including the need to address multiple pathophysiological mechanisms and the difficulty of accessing the affected tissues of the brain. Fibroblasts are a peripheral cell which can be obtained by a relatively simple biopsy directly from patients and expanded *in vitro* ^29^, making them useful in screening large numbers of compounds. Patient derived fibroblasts have been central in identifying and understanding mitochondrial dysfunction in primary mitochondrial diseases^11,12^ and importantly have been shown to recapitulate disease-related phenotypes in neurodegenerative diseases, exemplified in Parkinson’s disease^30,31^. High throughput screening in patient derived fibroblasts therefore has potential to facilitate drug discovery for modulators of the mitochondrial deficits observed in complex diseases such as LS and HD. Indeed fibroblasts have already been used successfully to identify compounds which have progressed into clinical trials and clinical use; Omaveloxalone, now approved for the treatment of Friedreich’s ataxia^32,33^, Ursodeoxycholic Acid, in phase 2 clinical trials for Parkinson’s disease^31,34,35^ and Sonlicrominal (KH176), now in phase 3 trials for primary mitochondrial diseases^36,37^ were identified as promising therapeutic compounds in initial testing undertaken in patient derived fibroblasts.

Here we evaluate disease relevant mitochondrial phenotypes in fibroblasts from patients with the childhood onset mitochondrial disease LS and the adult onset neurodegenerative disorder HD, and test selected compounds for their ability to alter the identified phenotypes. Since mitochondrial damage is already reported in fibroblasts from LS patients, we hypothesised that we would be able identify robust, consistent mitochondrial phenotypes across LS patient derived fibroblasts and would be able to identify small molecule modulators of this phenotype. We hypothesised that in HD cells, a more subtle mitochondrial phenotype would be present since HD is not primarily a mitochondrial disorder. Using high content imaging of the mitochondrial network, membrane potential and mitophagy flux, respirometry and measurements of cellular ATP, we identify patient specific phenotypes in LS and HD, including a novel deficit in mitophagy in LS cells. Our high content imaging profiling of tool compounds identifies A769662 as a modulator of the imbalance between functional and dysfunctional mitochondria found in both LS and HD cells. Our results find common mitochondrial dysfunction between otherwise unrelated diseases and highlight that screening for small molecule effects on mitochondrial phenotypes in patient derived fibroblasts could prove beneficial in finding novel treatments for neurological conditions.

## Materials and methods

### Patient derived fibroblasts

The following fibroblast cell lines were obtained from the NIGMS Human Genetic Cell Repository at the Coriell Institute for Medical Research: GM00969, GM28016, GM00498, GM20387, GM00409, GM28090, AG0222, GM00730, GM01169, GM23970, AG12956, GM04867, GM23248, GM06274. Details are shown in table 1. LS patient derived fibroblasts and their matched controls were cultured in a 50:50 mix of Eagle’s minimal medium (MEM, Corning):Dulbecco’s modified minimal essential medium (DMEM, Sigma) supplemented with 15% foetal bovine serum (Labtech), 100IU/ml penicillin, 100μg/ml streptomycin (sigma), 1mM sodium pyruvate (Sigma), 0.1mM amino acids (Thermofisher), 50μg/ml uridine (Sigma), and 1X minimum essential medium vitamins (Corning). HD patient derived fibroblasts and their matched controls were cultured in MEM with 10% foetal bovine serum, 100 IU/ml penicillin, 100μg/ml streptomycin (Thermofisher), 1mM sodium pyruvate, 0.1mM amino acids, 50μg/ml uridine, and 1X minimum essential medium vitamins. These culture conditions were used for all measurements except where galactose conditions are stated, in which case cells were cultured for the specified times in glucose free DMEM containing 10% foetal bovine serum, 100 IU/ml penicillin, 100μg/ml streptomycin, 1mM sodium pyruvate and 0.9 mg/ml galactose. Huntingtin gene repeat length in HD patients and corresponding controls was determined using repeat primed PCR sequencing.

**Table 1:** Cell lines obtained from Coriell for use in this study.

| **Fibroblast** | **Gender** | **Age at biopsy** | **Mutation / CAG repeat expansion** | **Coriell ID** |
| --- | --- | --- | --- | --- |
| Control 1 (LS) | Male | 3 years |  | GM00969 |
| LS1 | Male | 2 years | SURF1 paternally inherited C.370G>A (P.GLY124ARG) and maternally inherited C.751+1G>A | GM28016 |
| Control 2 (LS) | Female | 2 years |  | GM00498 |
| LS2 | Female | 6 years | SURF1 C.351T>G (P.TYR117*), C.312_321 DEL10INSAT | GM28037 |
| Control 3 (LS) | Male | 7 years |  | GM00409 |
| LS 3 | Female | 8 years | SURF1 (C.845_846DELCT) | GM28090 |
| Control 1 (HD)<br>(control for MMP and imaging phenotype; figures 8/9) | Male | 49 years | 15:ND | AG02222 |
| Control 1 (HD) | Female | 45 years | ND | GM00730 |
| HD1 | Male | 50 years | 17:44 | GM01169 |
| Control 2 (HD)<br>(control for MMP and imaging phenotype; figures 8/9) | Male | 50 years | 16:19 | GM23970 |
| Control 2 (HD) | Male | 50 years | ND | AG12956 |
| HD2 | Male | 49 years | 15:46 | GM04867 |
| Control 3 (HD) | Male | 55 years | 12:17 | GM23248 |
| HD3 | Female | 56 years | 16:43 | GM06274 |

### Immunoblotting

Cells were pelleted and lysed in radioimmunoprecipitation assay buffer (RIPA) buffer with the addition of phosphatase and protease inhibitors (Roche). Protein concentration was determined using Pierce BCA assay kit as per manufacturer’s instructions. 10 µg samples were separated using 7% (for probing HTT) or 12% sodium dodecyl sulfate polyacrylamide gel electrophoresis (Bio-Rad) and wet transferred to polyvinylidene difluoride membranes. Membranes were blocked in 5% milk (for OxPhos cocktail) or BSA. Primary antibodies were AMPK (2532S), p-AMPK (2531S), ULK1 (D8H5), p-ULK1 S555 (D1H4) (all cell signalling technologies), MTCO1 (ab14705), Ox-Phos cocktail (ab110411) (both Abcam), Huntingtin (MAB2166, sigma), GAPDH (60004-1-Ig, Proteintech). Secondary antibodies were horseradish peroxidase conjugated anti-rabbit (Dako) and anti-mouse IgG (sigma). Densitometry was carried out using enhanced chemiluminescence (Promega) on a Licor Odessey and processed using Image Studio 5.2.

### Mitochondrial function measurements

#### Complex Activity

Cells from confluent T75 cm^2^ flasks were pelleted and complex activity assays were carried out using a Complex IV Human Enzyme Activity Microplate Assay Kit (ab109909) as per the manufacturer’s instructions and previously published^31^ and measured on a PheraStar plate reader. Protein concentration was determined using Pierce BCA assay kit as per the manufacturer’s instructions and complex IV activity was normalised to protein content of each sample.

#### Total cellular ATP

Measurements of total cellular ATP measurements were carried out using ATPLite kit (Perkin Elmer) as described previously ^34^. Cells were plated at 5000 cells per well 72 hours before the assay in glucose containing media, which was changed to galactose media for galactose conditions. Luminescence measurements were made on an Omega plate reader (BMGLabtech). After ATP measurements, dsDNA levels were measured in the same lysates by incubating with 0.4X CyQUANT GR dye (Thermo Fisher Scientific) for 1 h at 37°C and measuring fluorescence emission at 520 nm on Omega plate reader. ATP levels were normalised to CyQuant values.

#### Respirometry

Cells were plated at a density determined for each pair of lines to allow 100% confluency on the day of the assay. Following optimisation of mitochondrial toxin concentrations, respiration levels were determined via the Mito Stress Test and measured using the Seahorse XFe analyzer (both Agilent) as per manufacturer’s instructions to measure oxygen consumption rate (OCR) in the absence and then presence of specific inhibitors. After measurement of basal OCR, 1.5µM of the ATP synthase inhibitor oligomycin was used to determine the proportion of OCR related to ATP production. 1µM (for HD cells) or 1.5µM (for LS cells). Carbonyl cyanide chlorophenylhydrazone (CCCP) was used to uncouple oxidative phosphorylation from ATP synthesis, causing cells to compensate for this by carrying out maximum levels of oxidative phosphorylation. Finally, complex I and complex III inhibitors, 0.5µM rotenone and 0.5µM antimycin A respectively, were used to inhibit all mitochondrial respiration and allowed us to determine the oxygen consumption specific to mitochondrial respiration. Cells were fixed with 3.7% paraformaldehyde after the assay and stained with 2 µM Hoechst (Sigma). Nuclei were imaged using an InCell Analzyer 2000 (GE Healthcare) and InCell Developer software used to count nuclei. OCR data are normalised throughout to number of cells.

#### Live fluorescent imaging

For live fluorescent imaging assays, cells were plated in 96 well plates in their respective culture medium. For each assay, cells were incubated with the relevant dyes in phenol free MEM for 1 h at 37°C. Dyes were removed and replaced with phenol red free media before imaging using a 40 x water objective on the Opera Phenix (Perkin Elmer). Ten fields of view were imaged per well, in 3 z planes. A minimum of 20 cells per technical replicate and therefore a minimum of 60 cells per cell line per experiment were analysed. Images were analysed using Harmony software (Perkin Elmer).

#### Mitochondrial membrane potential and morphology

72 or 48 hours prior to assaying, medium was replaced in the relevant wells with glucose free, galactose containing media. Mitochondria with a membrane potential were visualised using 80 nM tetramethylrhodamine (TMRM, sigma), the total mitochondrial population was visualised using 1 µM Mitotracker green (Thermo Fisher Scientific), and nuclei with 2 µM Hoechst (Sigma). Harmony software protocols were developed to segment nuclei, total mitochondria and mitochondria with membrane potential and to report morphology and intensity properties for each population. The following parameters are reported: total mitochondrial count, individual mitochondrial area, mitochondrial form factor, mitochondrial membrane potential, proportion of functional mitochondria, ratio of functional:total mitochondrial area and form factor.

#### Mitophagy

To investigate mitophagy in live cells, all mitochondria, mitochondria with a membrane potential, nuclei and lysosomes were visualised using 1 µM Mitotracker green (Thermo Fisher Scientific), 80 nM TMRM (Sigma), 2 µM Hoechst (Sigma) and 1 µM Lysotracker Deep Red (Thermo Fisher Scientific), respectively. Where specified, mitophagy was induced using 1µM Antimycin A (sigma) and 10µM Oligomycin (Sigma) immediately prior to imaging. Each field of view was imaged every hour for 6 hours. Harmony software protocols were developed to segment nuclei, mitochondria and lysosomes, to identify mitochondria which colocalise with lysosomes and to report morphology and intensity properties for each population as per those described in Schwartzentruber et al, 2020^38^.

#### Tool compound testing

Fibroblasts were plated as for live fluorescent imaging assays detailed above. At ∼70% confluency, cells were cultured in galactose containing medium for 48 hours before the addition of compounds or vehicle control, also in galactose containing medium, for a further 24 hours. UDCA (Sigma U5127) and Urolithin A (sigma, SML1791) were used at 1 μM and 100 nM, Mitochonic Acid (Sigma SML3166) was used at 10 μM and 5μM and A769662 (Sigma SML2578) was used at 100 μM and 10 μM. DMSO concentration for all conditions was 0.01%. Assays and analysis were carried out as per the mitochondrial membrane potential and morphology assay detailed above.

#### Statistical analysis

All statistical analyses were carried out on a minimum of three biological replicates and two technical replicates per biological replicate of each line. Unpaired t-test was used to determine significance between grouped control and patient derived fibroblasts. Data are represented as normalised to controls and hence the data are not normally distributed. Kruskal-Wallis with Dunn’s multiple comparisons were performed to determine significance between individual control and patient derived fibroblasts. Two-way ANOVAs were used to determine significance of media conditions or tool compound effects and any interaction effect between cell line and treatment. Multiple comparisons were corrected for using Šídák’s post hoc test. All statistical tests were carried out using GraphPad Prism software V10.4.1.

## Results

### In depth mitochondrial phenotyping of Leigh syndrome patient derived fibroblasts

#### Leigh syndrome patient derived fibroblasts display common complex IV deficits but donor-specific deficits in respiration

Since all three of the LS patients from which our fibroblasts are derived have mutations in the complex IV assembly factor SURF1, we first measured the protein levels of the mitochondrially encoded cytochrome C oxidase I (MTCO1), and the maximal activity of complex IV. Lower levels of MTCO1 did not reach significance in LS fibroblasts and lower maximal enzymatic activity of complex IV only reached significance in one LS fibroblast line compared to corresponding controls (figure 1A, 1B). We next measured total cellular ATP levels, mitochondrial membrane potential (MMP) and respiration in glucose and 48 or 72 hour galactose conditions to stimulate oxidative phosphorylation in the fibroblasts. Cellular ATP levels were not significantly different between control and LS cell lines, galactose conditions had little effect on ATP levels in either control or LS cells (figure 1C). Interestingly, under glucose-containing growth conditions, LS cells displayed significantly higher MMP than control cells, which was not sustained in 48 or 72 hour galactose conditions (figure 1D). Despite similar deficits in complex IV levels and activity between patients, we found that respiration deficits were specific to each individual donor-derived cell line. LS2 cells displayed strikingly reduced basal respiration, maximal respiration and spare respiratory capacity compared to their age-matched control cells. LS1 cells displayed significantly reduced basal respiration compared to their age-matched control, whilst deficits in other respiratory parameters do not reach significance and LS3 displayed no significant respiratory deficits compared to their age-matched control cells (figure 1E). Although LS3 displayed higher basal and induced extracellular acidification rate, no significant differences were observed in these parameters between control and LS cells, indicating no differences in glycolysis in LS (figure 1F).

**Figure 1:**
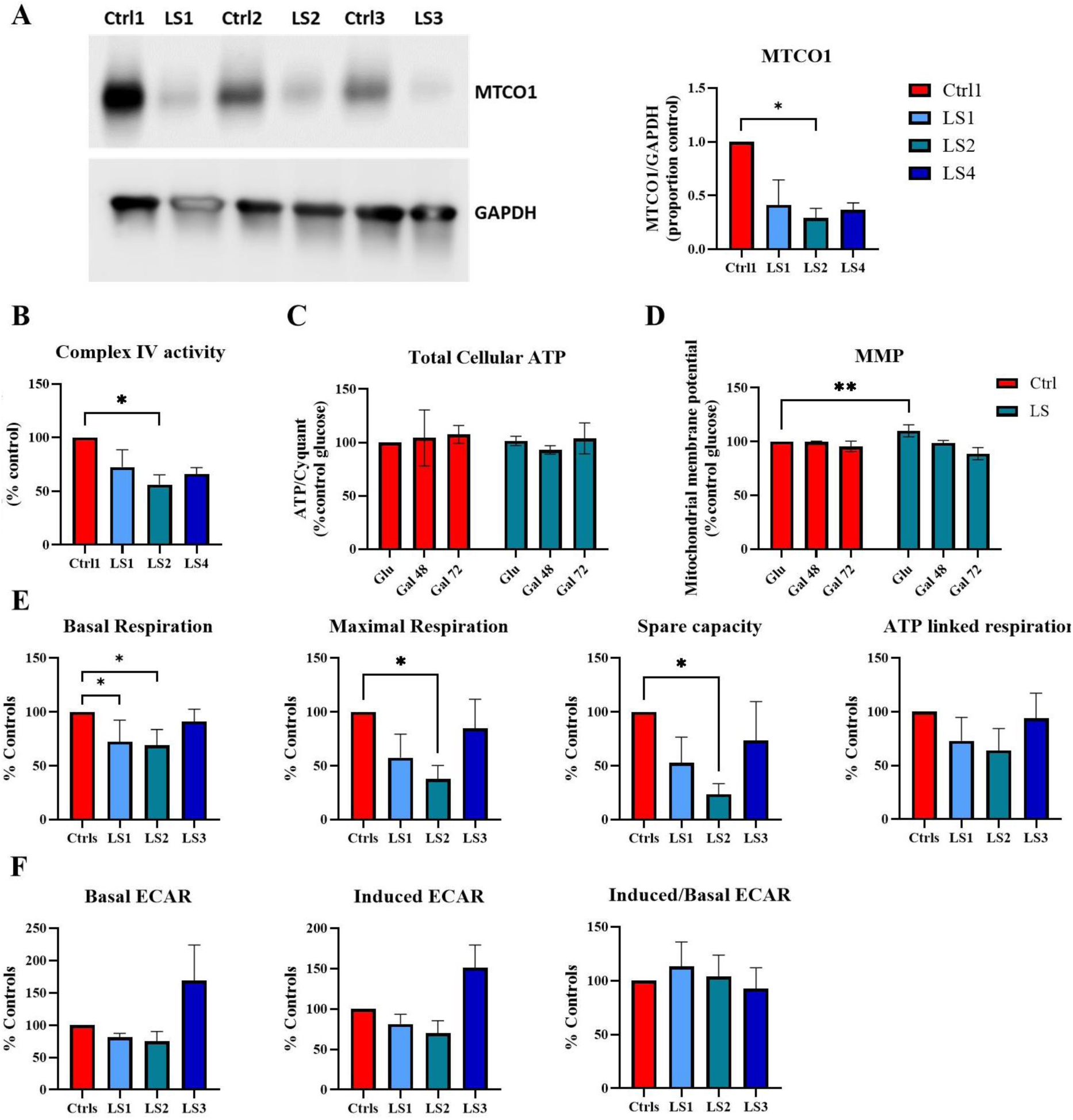
LS patient derived fibroblasts display damaged mitochondrial function. (A) Cytochrome c oxidase I (MTCO1) levels and quantification of MTCO1 levels normalised to GAPDH loading control, (B) Complex IV activity (C) total cellular ATP levels in glucose and 48 or 72 hour galactose conditions and (D) mitochondrial membrane potential in glucose and 48 or 72 hour galactose conditions in fibroblasts derived from three Leigh syndrome patients (LS, green) and three age matched control donors (Ctrl, red). Each data point represents the mean of three biological replicates from each individual cell line, normalised to age matched controls for each LS patient. (E) Basal respiration, maximal respiration, spare capacity and ATP linked respiration in fibroblasts derived from three Leigh syndrome patients (LS1, light blue, LS2, green, LS3, dark blue) and three age matched controls (Ctrls, red). (F) Basal, induced and basal/induced extracellular acidification rate in fibroblasts derived from three Leigh syndrome patients and three age matched controls (Ctrls, red). Bars represent the mean of three biological replicates for each cell line normalised to the age match control for each LS patient. Error bars represent standard deviation. C and D were analysed by two way ANOVA with Šídák’s multiple comparison test, A, B, E and F by Kruskal-Wallis with Dunn’s multiple comparisons, * = p<0.05.

#### High content imaging identifies a screenable phenotype in LS patient fibroblasts

In order to identify a phenotype in LS patient derived fibroblasts applicable to high throughput small molecule screening, we used high content imaging to assess parameters of mitochondrial morphology and function. As we observed in the bioenergetic assessment of LS patient derived fibroblasts, we identify a mitochondrial phenotype common to all three LS cell lines tested, as well as a subset of mitochondrial parameters which are specific to individual LS patient derived cell lines.

The proportion of mitochondria with a measurable membrane potential (proportion functional mitochondria) was significantly lower in all three LS cell lines compared to controls under 48 hours galactose conditions (figure 2A). The area of these functional mitochondrial compared to the total mitochondrial population was higher in LS patient cell lines compared to controls, though this only reached significance for LS2 at 72 hour galactose conditions (figure 2B). The functional network complexity compared to the total network complexity (indicated by functional form factor) was higher in LS1 and LS2 compared to their matched control in 48 hour galactose conditions (figure 2C). Taken together, these data indicate a functional mitochondrial network which consists of fewer, but larger and more fused functional mitochondria in LS cells compared to control cells. This assay is therefore suitable for identification of molecules aiming to decrease this maladaptive hyper fusion and increase mitochondrial fitness in patient cells.

**Figure 2:**
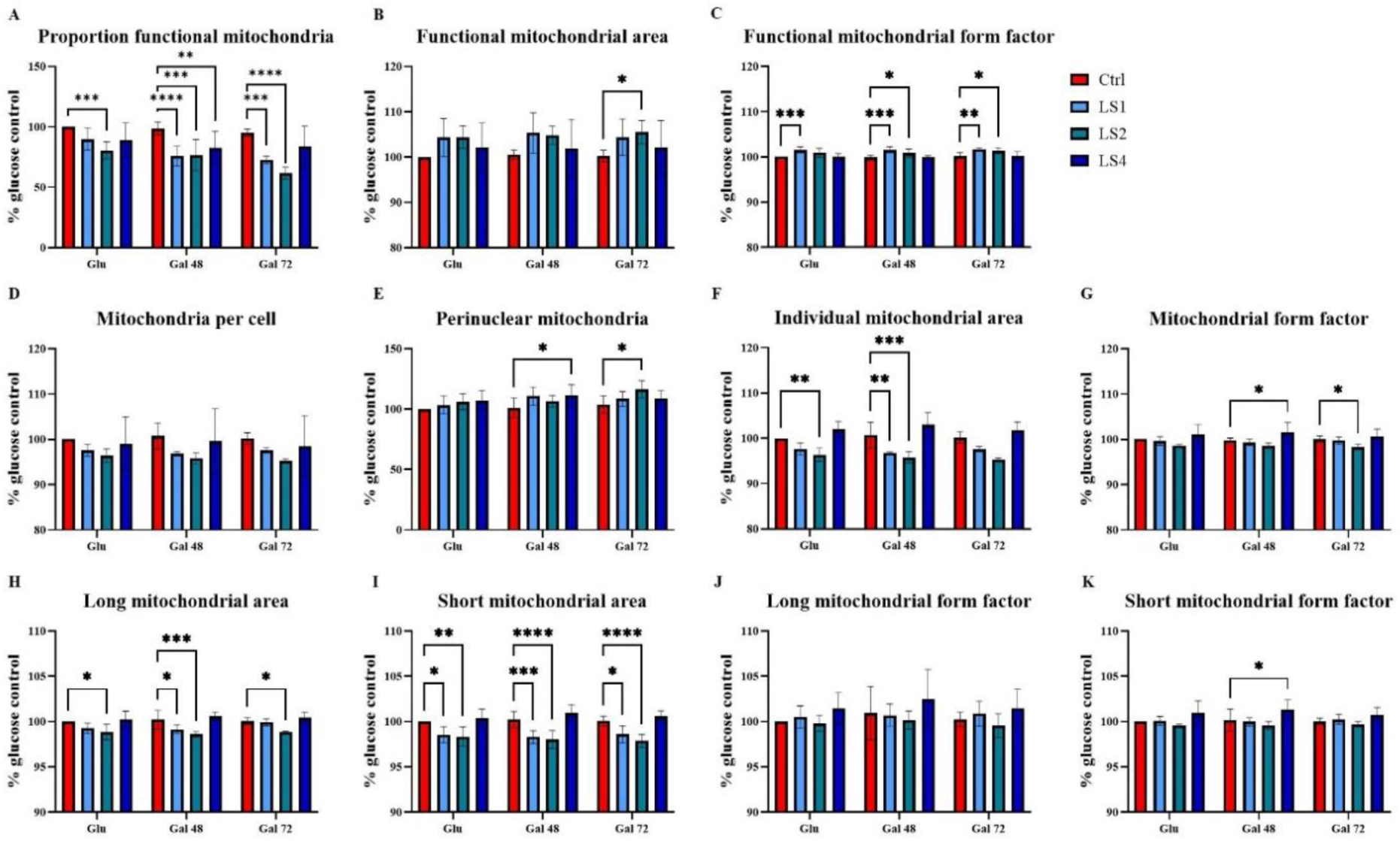
High content imaging of mitochondrial phenotype in LS patient derived fibroblasts. Mitochondrial morphology and functional ratios in LS patient derived fibroblasts (LS1, light blue, LS2 green, LS3 dark blue) compared to age matched control cells (Ctrls, red) in glucose or 48 or 72 hour galactose conditions. (A) Proportion of functional mitochondria is lower in all three LS patient derived fibroblasts in 48 hour galactose conditions compared to age matched controls. (B) Ratio of the average area of an individual mitochondrion with a membrane potential to the average area of an individual mitochondrion. (C) Ratio of the form factor of an individual mitochondrion with a membrane potential to the form factor of individual mitochondria. (D) Number of individual mitochondria per cell. (E) Number of mitochondria in the perinuclear space. (F) Individual mitochondrial area. (G) Mitochondrial form factor. (H) Area and (J) form factor of individual mitochondria in the long mitochondrial population. (I) Area and (K) form factor of individual mitochondria in the short mitochondrial population. Bars represent the mean of three biological replicates for each cell line normalised to the age match control for each LS patient. Error bars represent standard deviation. Data analysed by Two-way ANOVA with Šídák’s multiple comparisons test, * = p<0.05, ** = p<0.01, *** = p<0.001, **** = p<0.0001.

There were no significant differences in the total numbers of mitochondria between LS and healthy control fibroblasts. LS2 and LS3 displayed increased numbers of perinuclear mitochondria compared to controls under 72 and 48 hours galactose conditions respectively, indicating an altered distribution of mitochondria within the cell (figure 2D, 2E respectively). LS1 and LS2 cells had significantly smaller mitochondria under 48 hour galactose conditions and this is consistent in both long and short mitochondrial populations (figure 2F, 2H, 2I respectively).

Compared to glucose conditions, mitochondrial numbers per cell significantly increased in control cells in 72 hour galactose conditions (figure 3A). In control cells, MMP, proportion of functional mitochondria and perinuclear mitochondrial numbers did not change in the different growth conditions (figure 3B, 3C, 3D respectively). LS1 and LS2 cells also displayed significant increases in mitochondrial numbers under 72 hour galactose conditions, whilst LS3 did not. MMP and the proportion of functional mitochondria reduced in all three LS cell lines in 72 hour galactose conditions, indicating that compared to controls, LS cells are less able to maintain functional mitochondrial membrane potential under conditions which stimulate oxidative phosphorylation.

**Figure 3:**
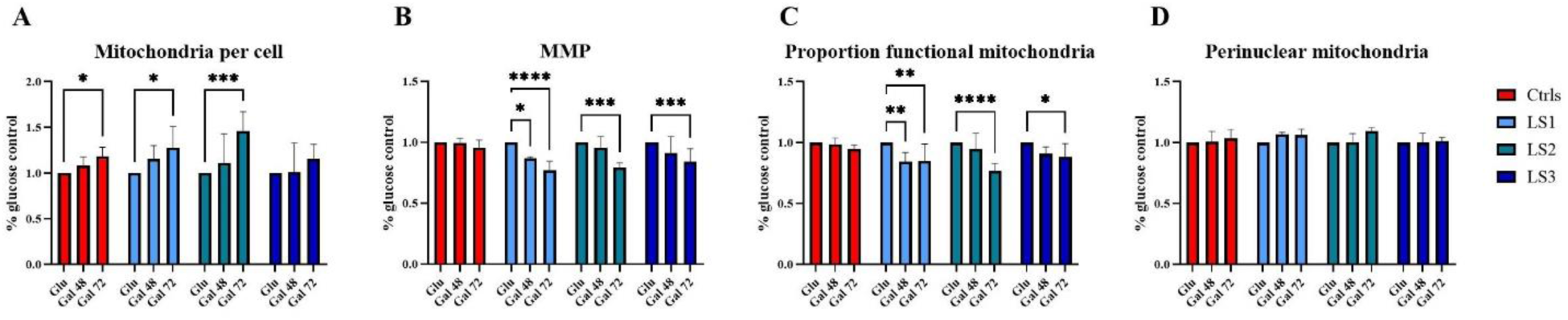
Mitochondrial membrane potential and proportion of functional mitochondria are damaged in galactose culture conditions in LS patient derived fibroblasts but not age matched control fibroblasts. (A) Mitochondria per cell increases under galactose (Gal) culture conditions in control (red), LS1 (light blue) and LS2 (green) fibroblasts but not in LS3 (dark blue) fibroblasts. (B) Mitochondrial membrane potential (MMP) and (C) the proportion of functional mitochondria are reduced in galactose culture conditions compared to glucose in all three LS cells but not in control cells. (D) Perinuclear mitochondrial numbers are not changed in galactose compared to glucose containing culture conditions. Data are normalised to the glucose values for each cell line, bars represent the mean of three biological replicates. Error bars represent standard deviation. Data analysed by Two-way ANOVA with Šídák’s multiple comparisons test, * = p<0.05, ** = p<0.01, *** = p<0.001, **** = p<0.0001.

#### Tool compound profiling identifies A769662 as an enhancer of proportion of functional mitochondria

To determine whether the phenotype we describe in LS patient cells using high content imaging can be rescued using small molecule compounds, we tested selected tool compounds using this imaging paradigm. Since we identified primary deficits in the proportion of mitochondria with functional membrane potential and network fusion in LS patient cells, we selected compounds reported to enhance mitochondrial function, membrane potential or the process of clearing dysfunctional mitochondria, mitophagy. Ursodeoxycholic acid (UDCA) is a bile acid reported to reduce mitochondrial induced apoptosis, reduce reactive oxygen species and enhance mitochondrial function although its direct mechanism of action is unclear^39^. A769662 is an activator of the metabolic regulator AMP-activated protein kinase (AMPK) found to rescue mitochondrial membrane potential^40^. Mitochonic acid 5 (MA5) and Urolithin A (UroA) are both reported activators of mitophagy^41,42^. These compounds were tested on mitochondrial phenotypes using concentrations reported to affect mitochondrial function and clearance in cell models by others^43–46^.

Neither UDCA, MA5 nor UroA had any significant effects on control or LS patient cells in the parameters we selected, although there was a trend towards reduced MMP in cells treated with MA5 and UroA (figure 4D). We found that 100μM A769662 significantly enhanced the proportion of functional mitochondria in both control and LS cells (figure 4E), increased the individual mitochondrial size in control cells (figure 4B) and decreased the number of perinuclear mitochondria in both LS and control cells (figure 4H). 100 μM A769662 also significantly reduced the ratio of the average individual functional mitochondrial area to the average overall individual mitochondrial area in control cells (figure 4F), likely indicating a removal of smaller, dysfunctional mitochondria. None of the compounds tested had a significant effect on mitochondrial form factor, membrane potential or functional/total mitochondrial form factor (figure 4C, 4D, 4G). These data confirm that the mitochondrial phenotype identified by high content imaging in LS patient fibroblasts is sensitive to identification of compound effects and suggests that A769662 has a beneficial effect on this phenotype. No compounds induced toxicity as measured by cell count (supplementary figure 1).

**Figure 4:**
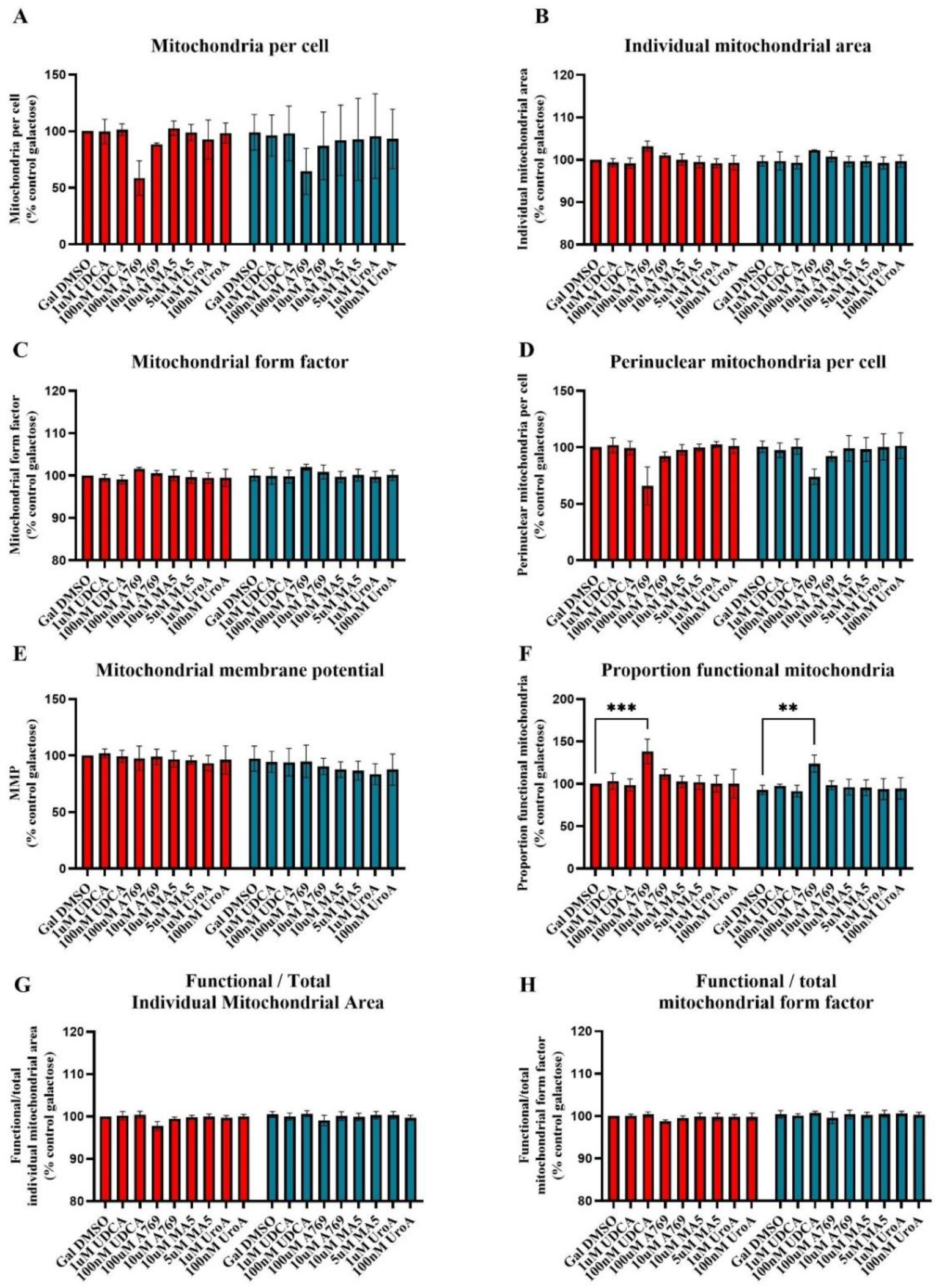
Small molecule profiling in LS patient derived fibroblasts (LS, green) compared to age matched control cells (Ctrls, red). Effect of 24 hour treatment with 1μM and 100nM UDCA, 100μM and 10μM A769662, 10 and 5 μM MA5 and 1 μM and 100nM UroA on (A) number of individual mitochondria per cell, (B) individual mitochondrial area (C) mitochondrial form factor, (D) MMP, (E) proportion of functional mitochondria (F) ratio of the average area of an individual mitochondrion with a membrane potential to the average area of individual mitochondria, (G) ratio of the form factor of an individual mitochondria with a membrane potential to the form factor of individual mitochondria and (H) the number of mitochondria in the perinuclear space. Bars represent the mean of three technical and three biological replicates for each cell line, normalised to vehicle (Gal DMSO) treated condition for the age matched control for each LS patient. Error bars represent standard deviation. Data analysed by Two-way ANOVA with Šídák’s multiple comparisons test, * = p<0.05, ** = p<0.01, *** = p<0.001, **** = p<0.0001.

#### A769662 acts via enhancing ULK1 phosphorylation and mitophagy in control and LS cells

Since A769662 is an allosteric activator of the metabolic regulator AMPK, which has been reported to activate mitophagy^47^, and since we identified a higher proportion of functional mitochondria and a trend toward reduced mitochondrial numbers, we hypothesised that A769662 acts via mitophagy to reduce the number of dysfunctional mitochondria and increase the proportion remaining mitochondria that are functional. ULK1 is a downstream target for phosphorylation by AMPK and plays a role in mitophagy^48–50^. We therefore sought to determine whether A769662 enhanced the phosphorylation of ULK1, indicating activation of AMPK. A769662 treatment significantly increased the levels or proportion of ULK1 phosphorylation in control, LS1 and LS3 cells (figure 5A, B).

**Figure 5:**
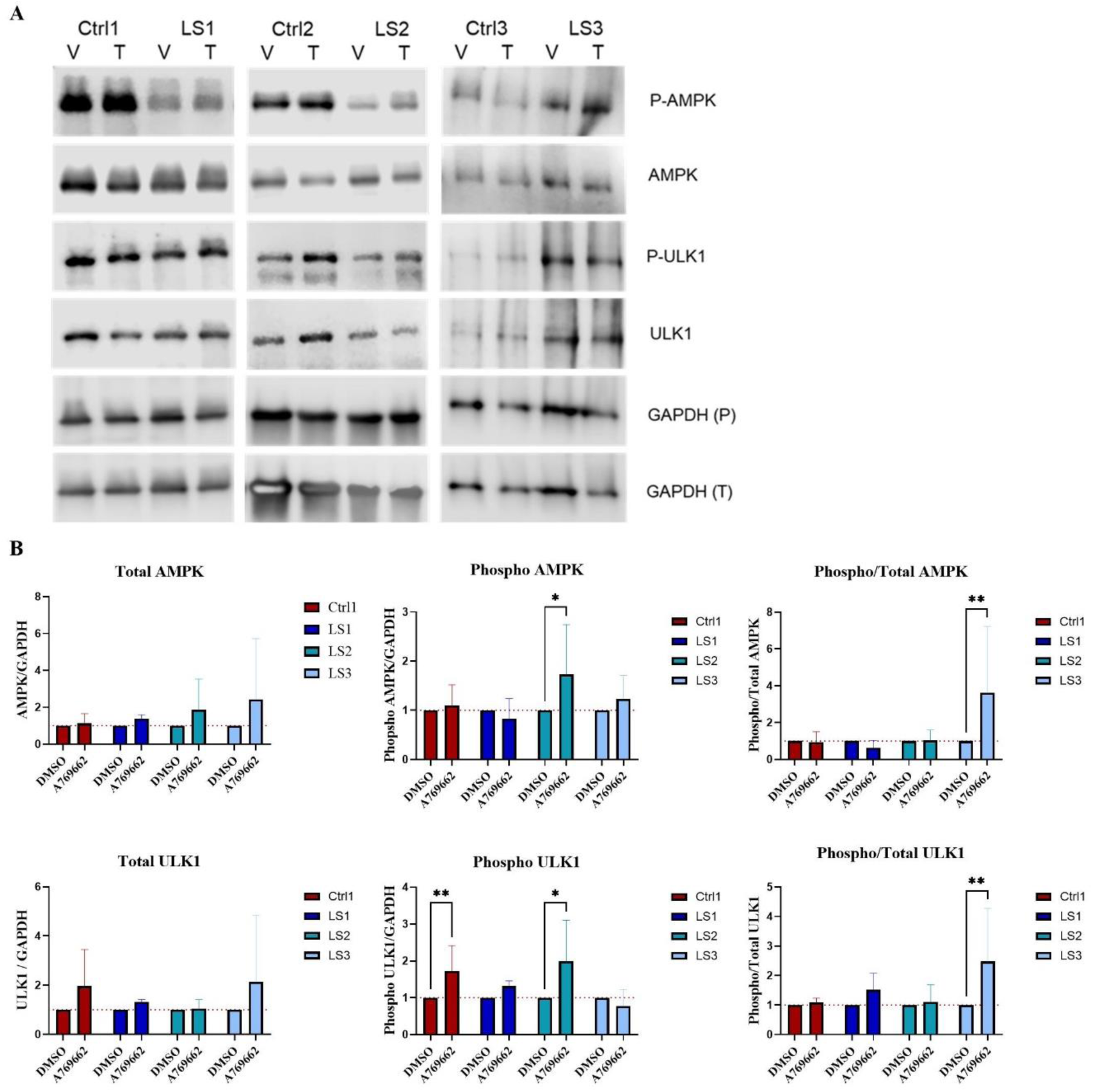
Effect of A769662 on AMPK and ULK1 phosphorylation. (A) Levels of AMPK, phosphorylated AMPK, ULK1, phosphorylated ULK1 and housekeeping protein for total AMPK/ULK1 blots GAPDH (T), and for phosphorylated AMPK/ULK1 blots GAPDH (P), in control and LS cells treated for 24 hours with DMSO vehicle (V) or 100 μM A769662 (A769) treatment (T). (B) Quantification of (A). Total, phosphorylated, and ratio of phosphorylated / total AMPK (top graphs) and ULK1 (bottom graphs) levels normalised to GAPDH and DMSO treatment in LS cells (blues) compared to control cells (red). Values represent three biological replicates for each cell line and treatment. Error bars represent standard deviation. Data analysed by One way ANOVA with Holm-Šídák’s multiple comparisons test, * = p<0.05, ** = p<0.01, p=<0.0001.

To determine whether A769662 enhanced mitophagy, we measured the percentage of mitochondria colocalised with lysosomes after 24 hours of A769662 treatment under basal glucose conditions, and immediately after the induction of mitophagy with 1 μM Oligomycin and 4 μM Antimycin A, for a period of 6 hours. We detected a small but significant basal deficit in mitophagy levels in LS cells compared to age matched controls (figure 6B) which has not been reported before. 1 μM A769662 treatment reduced the percentage of mitochondria at lysosomes in basal conditions however no effect was seen with higher concentrations (figure 6D, F). We identified a significantly smaller percentage of mitochondria colocalising to lysosomes upon mitophagy stimulation in LS compared to control cells over the 6 hour period (figure 6C). 100 μM A769662 significantly enhanced mitophagy in LS cells (figure E), driven by an immediate increase in mitochondria colocalising with lysosomes upon mitophagy induction, indicated by assessment of the first time point in our time lapse experiments (figure 6G).

**Figure 6:**
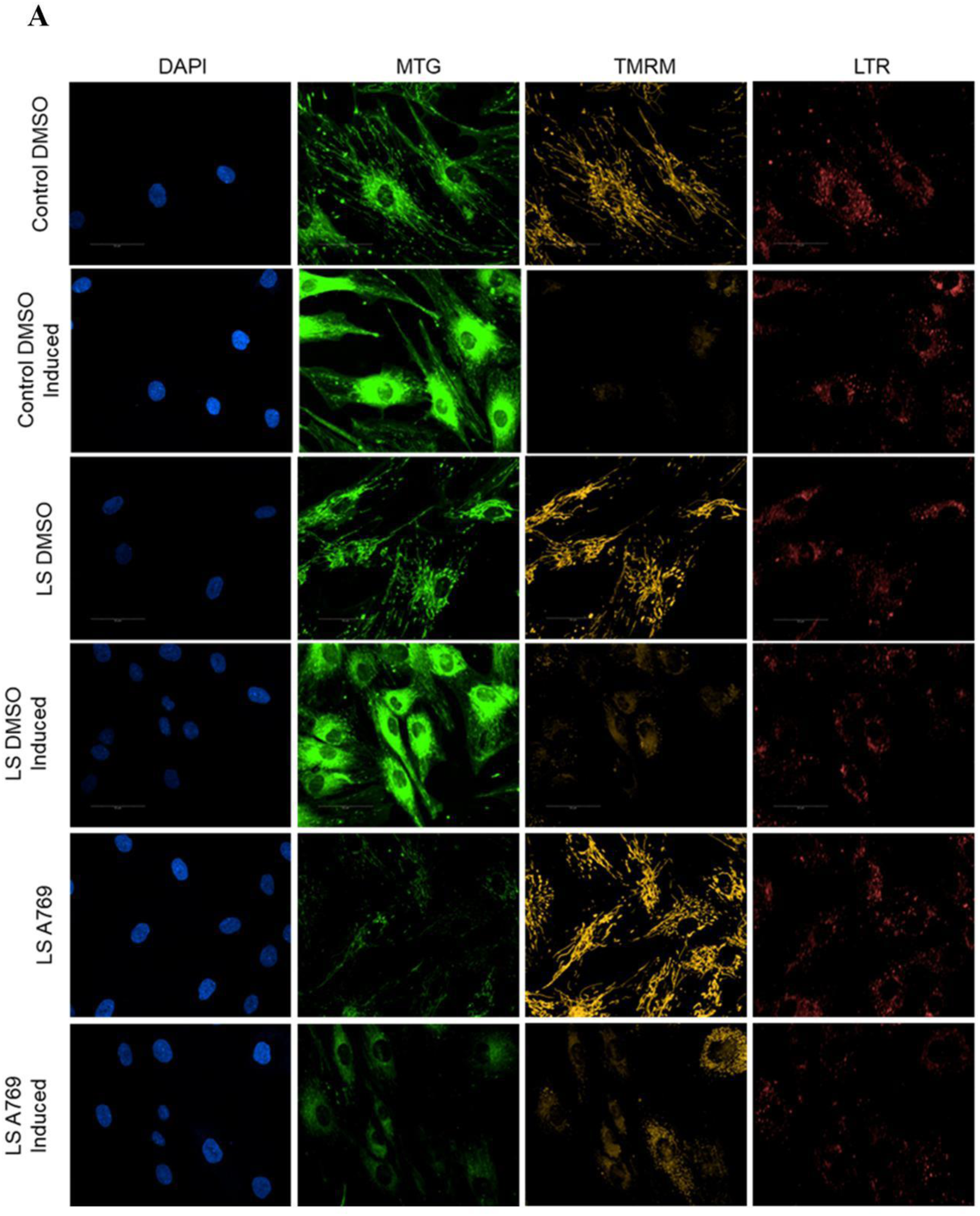
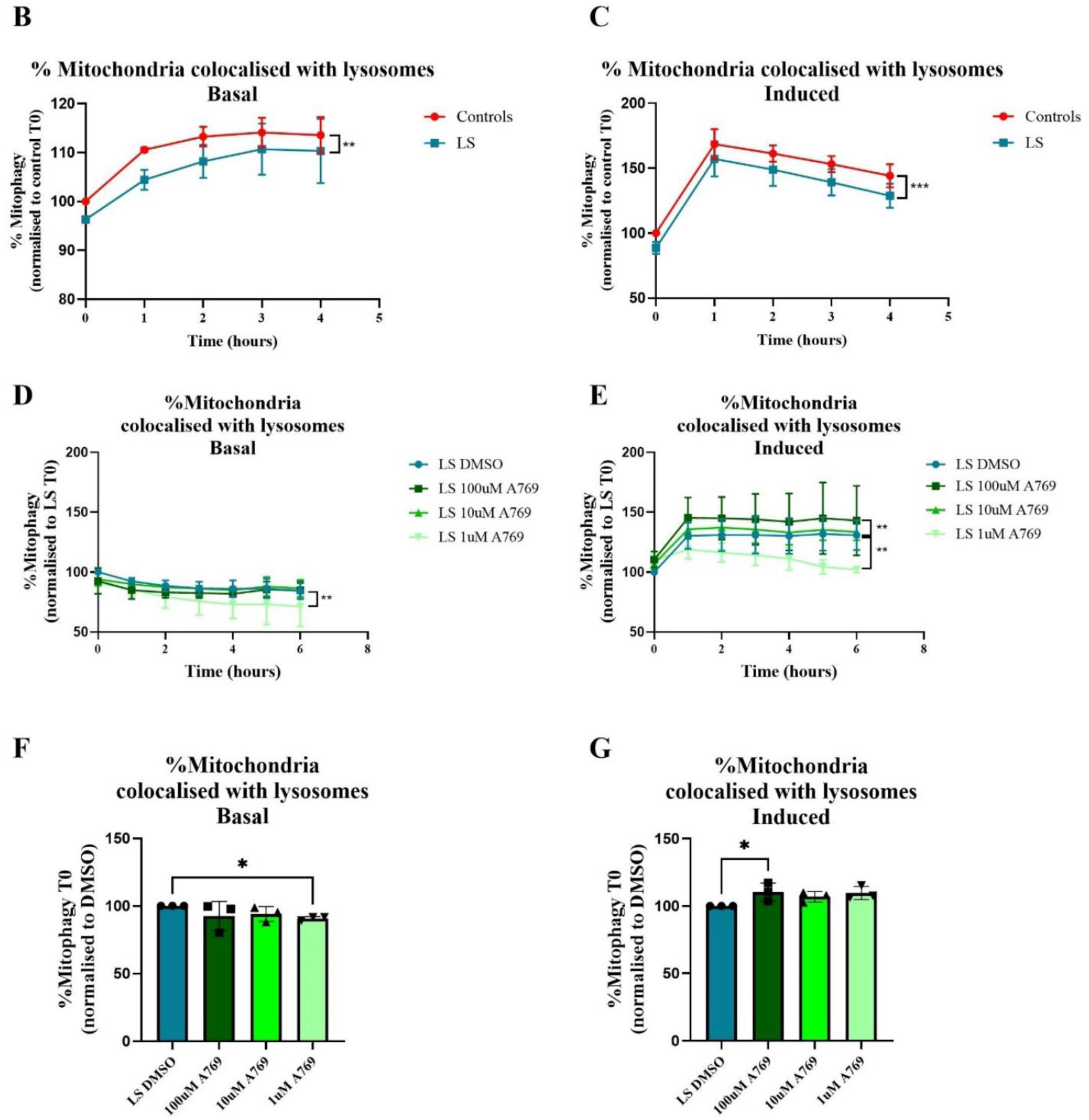
A769662 enhances induced mitophagy in LS patient derived fibroblasts. (A) Representative images of nuclei (DAPI), total mitochondrial population (Mitotracker green, MTG), mitochondrial membrane potential (TMRM), and lysosomes (Lysotracker deep red, LTR) from live mitophagy assay after 24 hour treatment with DMSO vehicle or 100 μM A769662, in basal conditions and immediately after the addition of 1 μM Oligomycin and 4 μM Antimycin A (Induced conditions). (B-E) Mitophagy, calculated as the percentage of total mitochondria which colocalised with lysosomes. Mitophagy over 6 hours in control and LS cells in basal (B) and induced (C) conditions, normalised to control cell lines at T0. Mitophagy over 6 hours in LS cells pre-treated for 24 hours with DMSO or 100, 10, 1 μM A769662 in basal (D) and induced (E) conditions, normalised to T0 of the DMSO vehicle treated condition for each LS patient. Timepoint 0 plotted for basal (F) and induced (G) conditions. Lines and bars represent the mean of three technical and three biological replicates for each cell line. Error bars represent standard deviation. Data analysed by Two-way ANOVA, with Holm-Šídák’s multiple comparisons test for treatment effects, * = p<0.05, ** = p<0.01, *** = p<0.001, **** = p<0.0001.

### In depth mitochondrial phenotyping of Huntington’s Disease patient derived fibroblasts

#### Bioenergetic deficiencies in HD patient fibroblasts

Having confirmed the capability of high content image screening to identify small molecule modulators of mitochondrial phenotypes for LS, a condition caused by a primary mitochondrial deficit, we next sought to apply this technique to HD. Mitochondrial dysfunction is implicated in HD, but the genetic cause of HD is not a mitochondrial genetic defect, as in LS. Previous work in HD describes mitochondrial dysfunction in various HD models, including evidence of morphological changes and respiratory complex damage in patient fibroblasts^19,20,26^. HTT is expressed at low levels in fibroblasts and we therefore first confirmed the presence of HTT at low levels in both control and patient derived fibroblasts (figure 7A).

**Figure 7:**
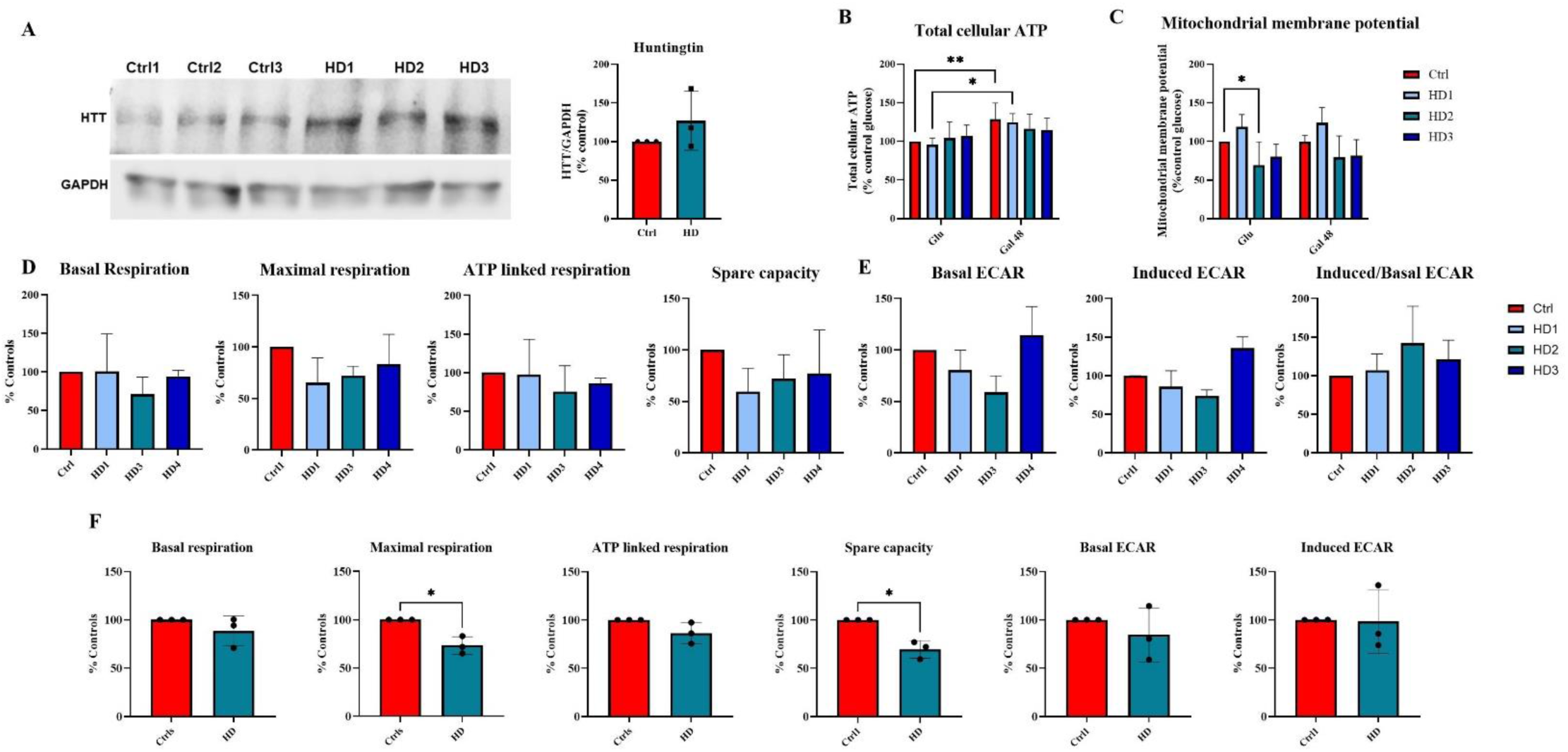
HD patient derived fibroblasts display damaged mitochondrial function. (A) HTT protein levels and quantification normalised to GAPDH, (B) total cellular ATP levels in glucose and 48 hour galactose conditions and (C) mitochondrial membrane potential in glucose and 48 hour galactose conditions in fibroblasts derived from three Huntington patients (HD1, light blue, HD2, green, HD3, dark blue) and three age matched control donors (Ctrl, combined in red bar). Each data point represents the mean of three biological replicates from each individual cell line, normalised to age matched control for each HD patient. (D) Basal respiration, maximal respiration, spare capacity and ATP linked respiration in fibroblasts derived from three HD patients (HD1, light blue, HD2, green, HD3, dark blue) and three age matched controls (Ctrls, red). (E) Basal, induced and basal/induced extracellular acidification rate in fibroblasts derived from three HD patients (HD1, light blue, HD2, green, HD3, dark blue) and three age matched controls (Ctrls, red). Bars represent the mean of three biological replicates for each cell line normalised to the age matched control for each HD patient. (F) Parameters in D and E with data for patients combined so that each data point represents the mean of three biological replicates for a single patient and the bar represents the mean for all three patients. Error bars represent standard deviation. Data analysed by Wilcoxon (A, F) and two-way ANOVA with Šídák’s multiple comparisons test (B-D) * = p<0.05, ** = p<0.01.

Since mitochondrial dysfunction is implicated in HD pathogenesis, but the underlying genetic cause is not mitochondrial, we hypothesised that bioenergetic deficiencies would be present in HD patient derived fibroblasts, but that the mitochondrial phenotype may vary between individuals. As in LS cells, total amounts of cellular ATP were not significantly different between the control and HD cell lines. However, 48 hour galactose conditions induced increased ATP levels in the control cells and in HD1 cells, indicating their shift from glycolysis to oxidative phosphorylation which yields more ATP. HD2 and HD3 did not display this increase (figure 7B). HD2 was found to have significantly lower MMP compared to controls in glucose containing conditions, whilst there were no differences between controls and HD1 and HD3, and no differences between control and any HD cells after 48 hours of culture in galactose (figure 7C). There were no significant differences in respiration rates or extracellular acidification rates between control cells and any individual HD cell line (figure 7D).

#### Mitochondrial morphology is altered in fibroblasts from patients with Huntington’s disease

Having identified functional mitochondrial parameters that were damaged in HD cells, we again sought to determine whether a robust mitochondrial phenotype could be identified using high content imaging. As anticipated, we found individual differences in mitochondrial morphology and function between patient derived cell lines. The proportion of mitochondria with a membrane potential appears lower in both HD2 and HD3 compared to controls but this only reaches significance in HD2 in glucose and 48 hour galactose conditions (figure 8A). HD2 and HD3 both have smaller functional mitochondria as compared to their total mitochondrial population, in glucose and 72 hour galactose conditions respectively (figure 8B) but no changes in the functional network fusion were found in HD cells compared to controls (figure 8C).

**Figure 8:**
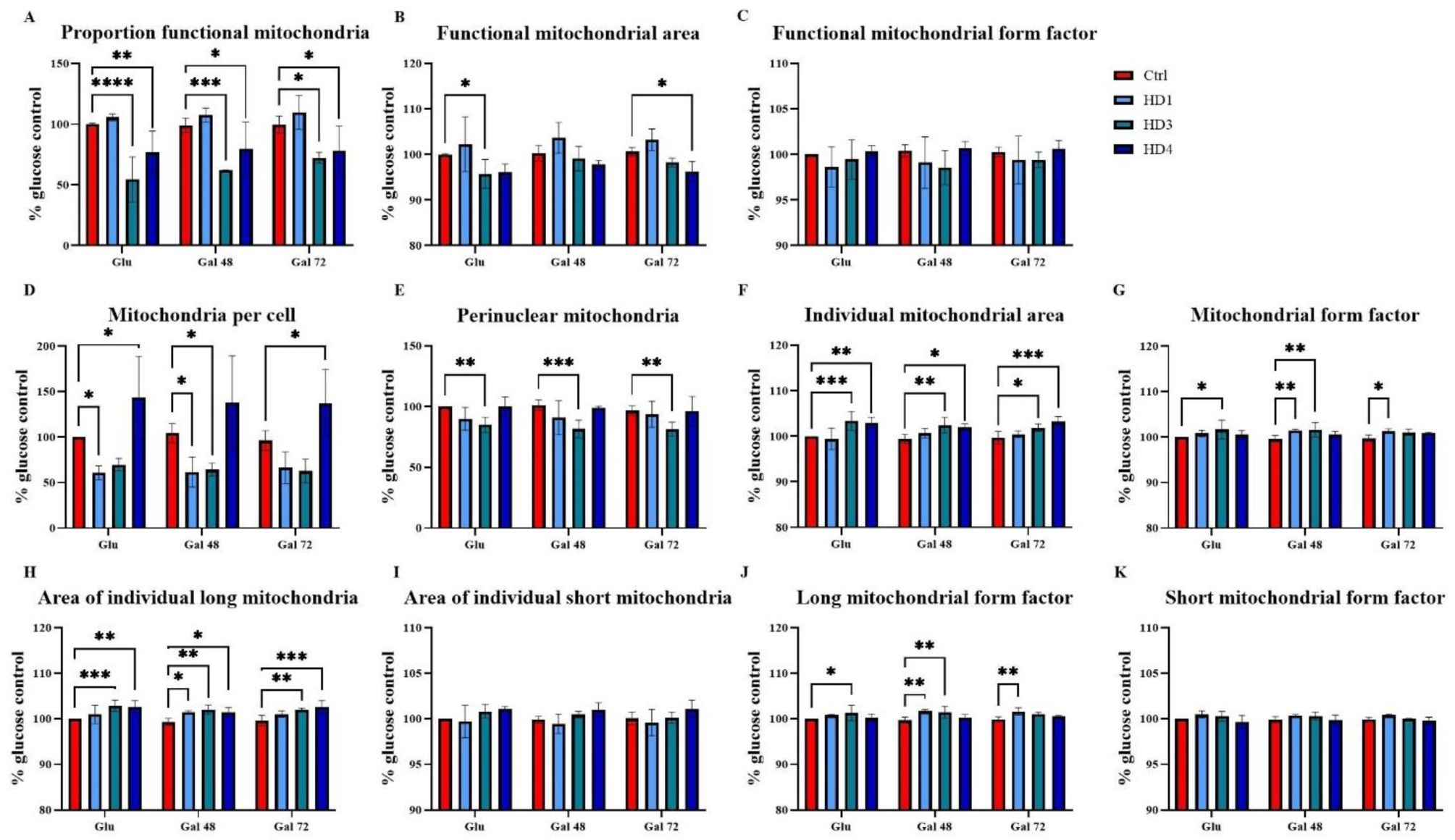
Mitochondrial morphology and functional ratios in HD patient derived fibroblasts. (HD1, light blue, HD2 green, HD3 dark blue) compared to control cells (Ctrls, red) in glucose or 48 or 72 hour galactose conditions. (A) Proportion of functional mitochondria, (B) ratio of the average area of an individual mitochondrion with a membrane potential to the average area of individual mitochondria, (C) ratio of the form factor of an individual mitochondria with a membrane potential to the form factor of individual mitochondria, (D) number of individual mitochondria per cell, (E) number of mitochondria in the perinuclear space, (F) individual mitochondrial area, (G) mitochondrial form factor, (H) area and (J) form factor of individual mitochondria in the long mitochondrial population and (I) area and (K) form factor of individual mitochondria in the short mitochondrial population. Bars represent the mean of three biological replicates for each cell line normalised to the combined mean of multiple controls. Error bars represent standard deviation. Data analysed by Two-way ANOVA with Šídák’s multiple comparisons test, * = p<0.05, ** = p<0.01, *** = p<0.001, **** = p<0.0001..

HD1 and HD2 have fewer mitochondria than control cells, reaching significance in different media conditions. Perinuclear mitochondria count was also lower in HD2 in 48 hour galactose conditions compared to control cells, whilst the number of mitochondria in HD3 was higher than control cells under glucose and 72 hour galactose conditions (figures 8D and E). Larger individual mitochondria found in HD2 and HD3 compared to control cell lines, and increased form factor in HD1 and HD2 cell lines compared to controls, indicates an elongation or a fusion of the mitochondrial network in HD fibroblasts (figures 8F and G). Increased area of mitochondria in the long mitochondrial population, in all HD cell lines compared to controls in 48 hour galactose conditions, and increased form factor of this population in HD1 and HD2 compared to controls support this phenotype (figures 8H and J). There were no significant differences in the population of short mitochondria between patient and control cells (figures 8I and K).

Taken together, these data indicate that all HD cell lines have enlarged mitochondria in a fused network but that functional damage manifests differently across individual cell lines; HD1 cells have less mitochondria than control cells but which appear to retain membrane potential, HD2 cells have less mitochondria which are also functionally damaged, and HD3 cells have more mitochondria but their functional morphology is damaged.

We find no significant difference in the number of mitochondria, MMP, proportion of functional mitochondria or number of perinuclear mitochondria in control cells grown in glucose compared to control cells grown in galactose for 48 or 72 hours. The same is true for HD1 and HD3, with no significant differences between growth media conditions. HD2 cells however, which have lower proportion of functional mitochondria compared to control cells, display an increase in both MMP and proportion of functional mitochondria after 72 hours of culture in galactose containing media (figure 9).

**Figure 9:**
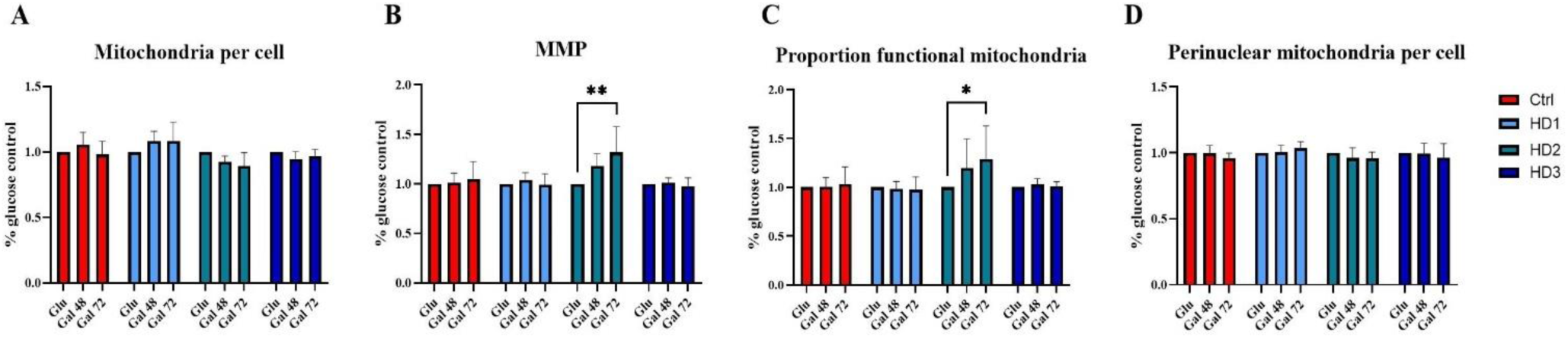
Effect of 48 and 72 hour galactose culture conditions (48 Gal and 72 Gal respectively) on HD patient derived fibroblasts. (HD1 light blue, HD2 green, HD3 dark blue) and age matched control fibroblasts (Ctrl, red). (A) Mitochondria per cell, (B) mitochondrial membrane potential (MMP), (C) the proportion of functional mitochondria and (D) perinuclear mitochondrial number. Data are normalised to the glucose values for each cell line, bars represent the mean of three biological replicates. Error bars represent standard deviation. Data analysed by Two way ANOVA with Šídák’s multiple comparisons test, * = p<0.05, ** = p<0.01.

#### Tool compound profiling identifies small molecule effects in HD cells

We next used the same selected tool compounds that we tested in LS cells to determine if their effects are also detectable in our HD cells in which we detect altered mitochondrial network morphology as well as phenotypes specific to each individual. Consistent with our findings in LS cells, we found that 100 μM A769662 enhanced the proportion of functional mitochondria in both control and HD cells (figure 10F) with little effect on area and form factor measurements (figure 10B and C) and a similar trend towards a reduction in overall mitochondria per cell (figure 10A). We saw no significant effects of the other compounds tested, although there was a trend toward Mitochonic acid 5 and Urolithin A reducing MMP which is consistent with their reported mode of action (figure 10E). No compounds induced toxicity as measured by cell count (supplementary figure 2).

**Figure 10:**
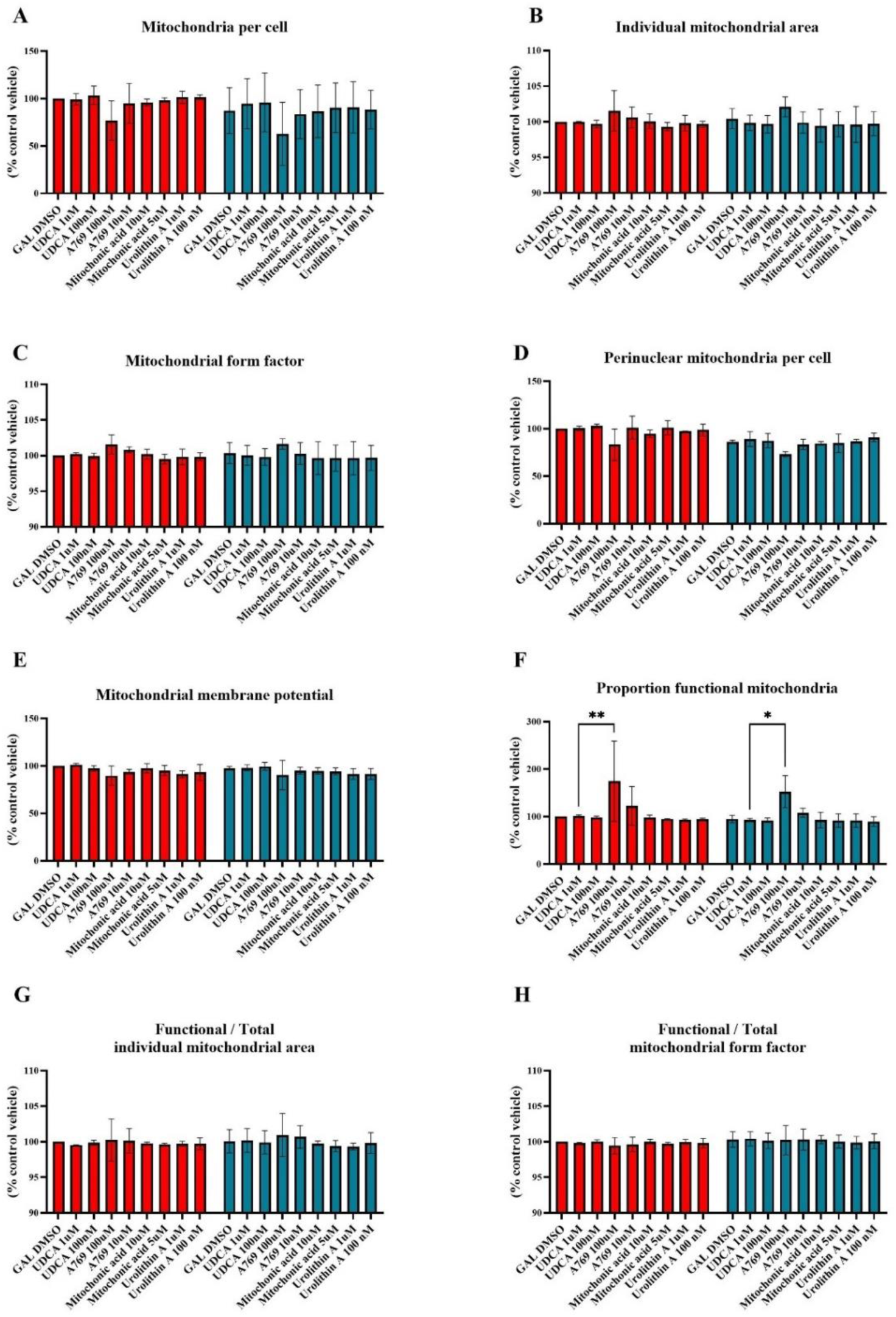
Small molecule assessment in HD patient derived fibroblasts. (HD, green) compared to age matched control cells (Ctrls, red). Effect of 24 hour treatment with 1 μM and 100 nM UDCA, 100 μM and 10 μM A769662, 10 and 5 μM MA5 and 1 μM and 100 nM UroA on (A) number of individual mitochondria per cell, (B) individual mitochondrial area (C) mitochondrial form factor, (D) MMP, (E) proportion of functional mitochondria (F) ratio of the average area of an individual mitochondria with a membrane potential to the average area of individual mitochondria, (G) ratio of the form factor of an individual mitochondria with a membrane potential to the form factor of individual mitochondria and (H) the number of mitochondria in the perinuclear space. Bars represent the mean of three technical and three biological replicates for each cell line, normalised to vehicle (Gal DMSO) treated condition for the age matched control for each HD patient. Error bars represent standard deviation. Data analysed by Two-way ANOVA with Šídák’s multiple comparisons test, * = p<0.05, ** = p<0.01.

#### A769662 does not act via enhancing mitophagy in older control and HD cells

As in LS cells, we next sought to determine whether A769662 enhanced ULK1 phosphorylation and mitophagy in HD cells to reduce the number of dysfunctional mitochondria and increase the proportion remaining that are functional. A769662 treatment significantly increased the levels or proportion of total or phosphorylated ULK1 control and HD cells (figure 11A, B).

**Figure 11:**
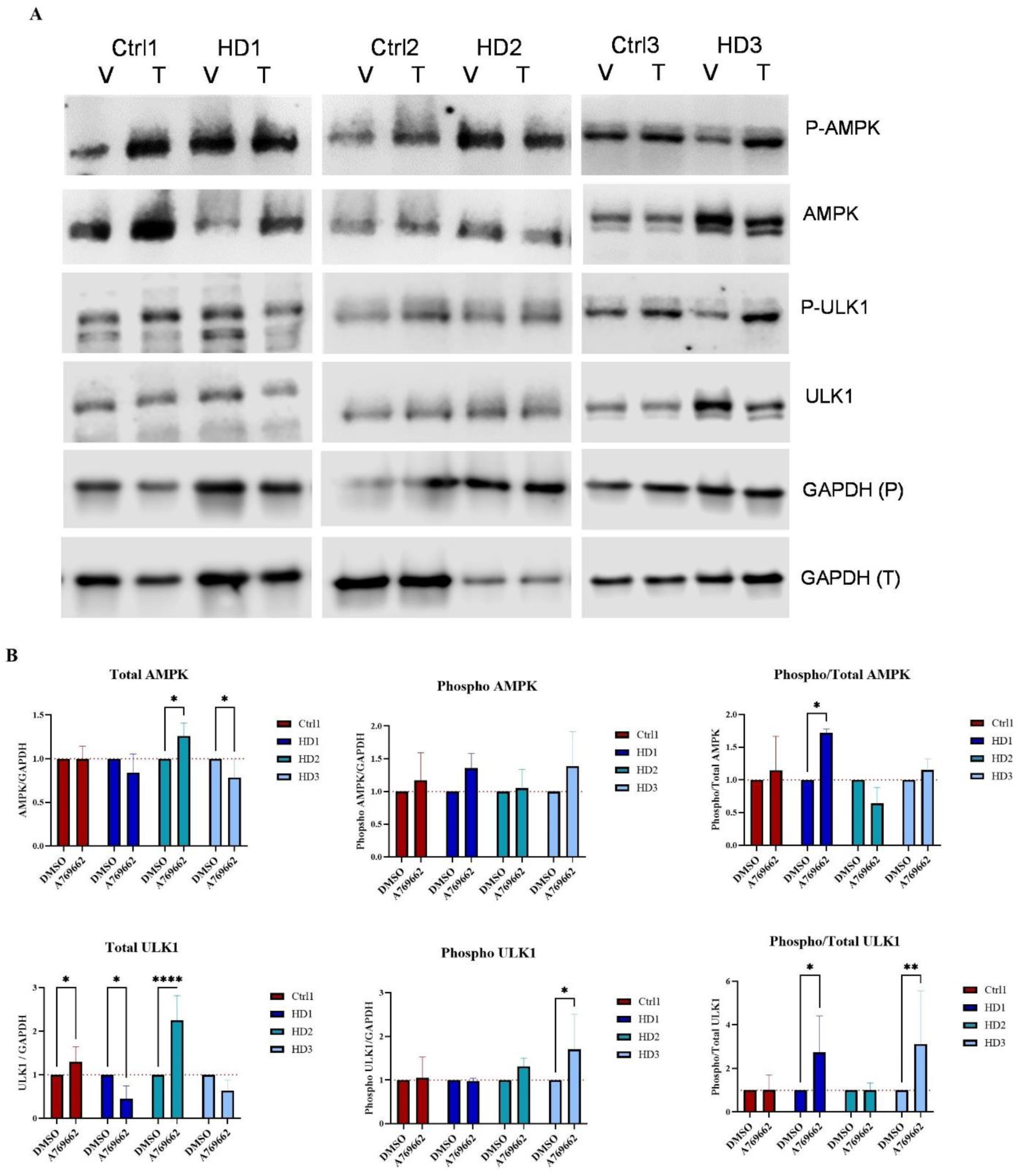
Effect of A769662 on AMPK and ULK1 phosphorylation. (A) Levels of AMPK, phosphorylated AMPK, ULK1, phosphorylated ULK1 and housekeeping protein for total AMPK/ULK1 blots GAPDH (T), and for phosphorylated AMPK/ULK1 blots GAPDH (P), in control and HD cells treated for 24 hours with DMSO vehicle (V) or 100 μM A769662 treatment (T). (B) Quantification of (A). Total, phosphorylated, and ratio of phosphorylated / total AMPK (top graphs) and ULK1 (bottom graphs) levels normalised to GAPDH and DMSO treatment in HD cells (blues) compared to control cells (red). Values represent three biological replicates for each cell line and treatment. Error bars represent standard deviation. Data analysed by One way ANOVA with Holm-Šídák’s multiple comparisons test, * = p<0.05, ** = p<0.01, p=<0.0001.

Under both basal (figure 12B) and mitophagy inducing conditions (1μM Oligomycin and 4μM Antimycin A, figure 12C) we found higher percentages of mitochondria colocalising with lysosomes in HD cells compared to age matched controls across the period of 6 hours, indicating either more mitochondria undergoing mitophagy, or a stalling of mitophagy at the point of the lysosomes, leading to a build-up of mitochondria in lysosomes. To determine whether A769662 enhanced mitophagy in these cells, we measured the percentage of mitochondria colocalised with lysosomes after 24 hours of A769662 treatment under basal conditions, and immediately after the induction of mitophagy for a period of 6 hours. In older control and HD cell lines, A769662 treatment did not enhance the percentage of mitochondria at lysosomes under basal or mitophagy inducing conditions (figure 12D-G). Since the effect of A769662 on enhancing the proportion of functional mitochondria was consistent across all patient and control cell lines, these data suggest that, although this effect may be in part mediated by the role of ULK1 activation on mitophagy, particularly in younger cells and LS cells, it is likely that other AMPK related mechanisms are involved in addition.

**Figure 12:**
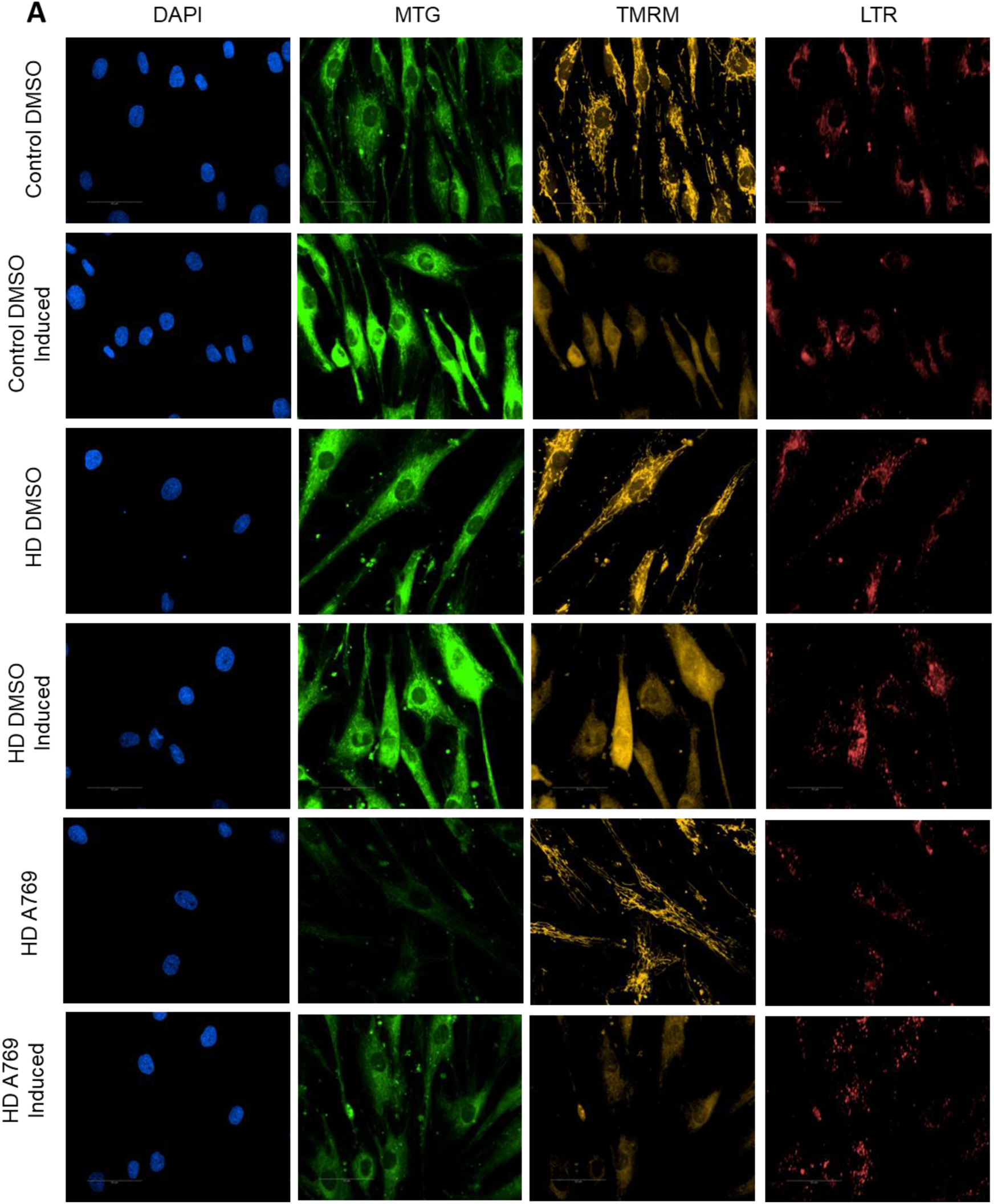
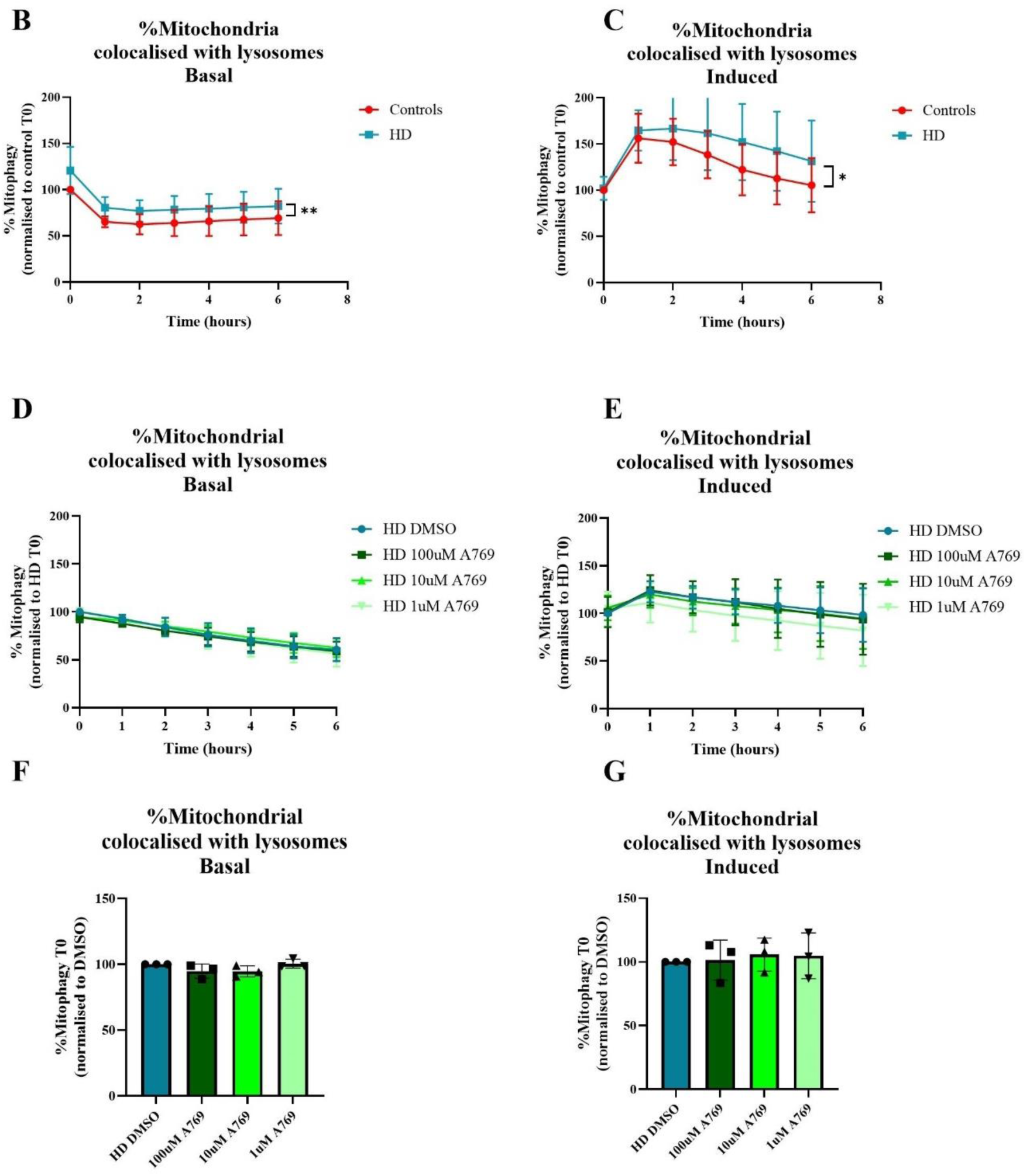
A769662 does not alter mitophagy in HD patient derived fibroblasts. (A) Representative images of nuclei (DAPI), total mitochondrial population (mitotracker green, MTG), mitochondrial membrane potential (TMRM), and lysosomes (lysotracker deep red, LTR) from live mitophagy assay after 24 hour treatment with DMSO vehicle or 100 μM A769662, in basal conditions and immediately after the addition of 1 μM Oligomycin and 4 μM Antimycin A (Induced). (B-G) Mitophagy, calculated as the percentage of total mitochondria which colocalise with lysosomes. Mitophagy over 6 hours in control and HD cells in basal (B) and induced (C) conditions normalised to time point 0 (T0) for each control cell line. Mitophagy over 6 hours in HD cells pre-treated for 24 hours with DMSO or 100, 10, 1 μM A769662 in basal (D) and induced (E) conditions, normalised to the DMSO vehicle condition at T0. T0 data from D and E shown in F and G respectively. Lines and bars represent the mean of three technical and three biological replicates for each cell line. Error bars represent standard deviation. Data analysed by Two-way ANOVA with Holm-Šídák’s multiple comparisons test, * = p<0.05, ** = p<0.01.

**Figure 13:**
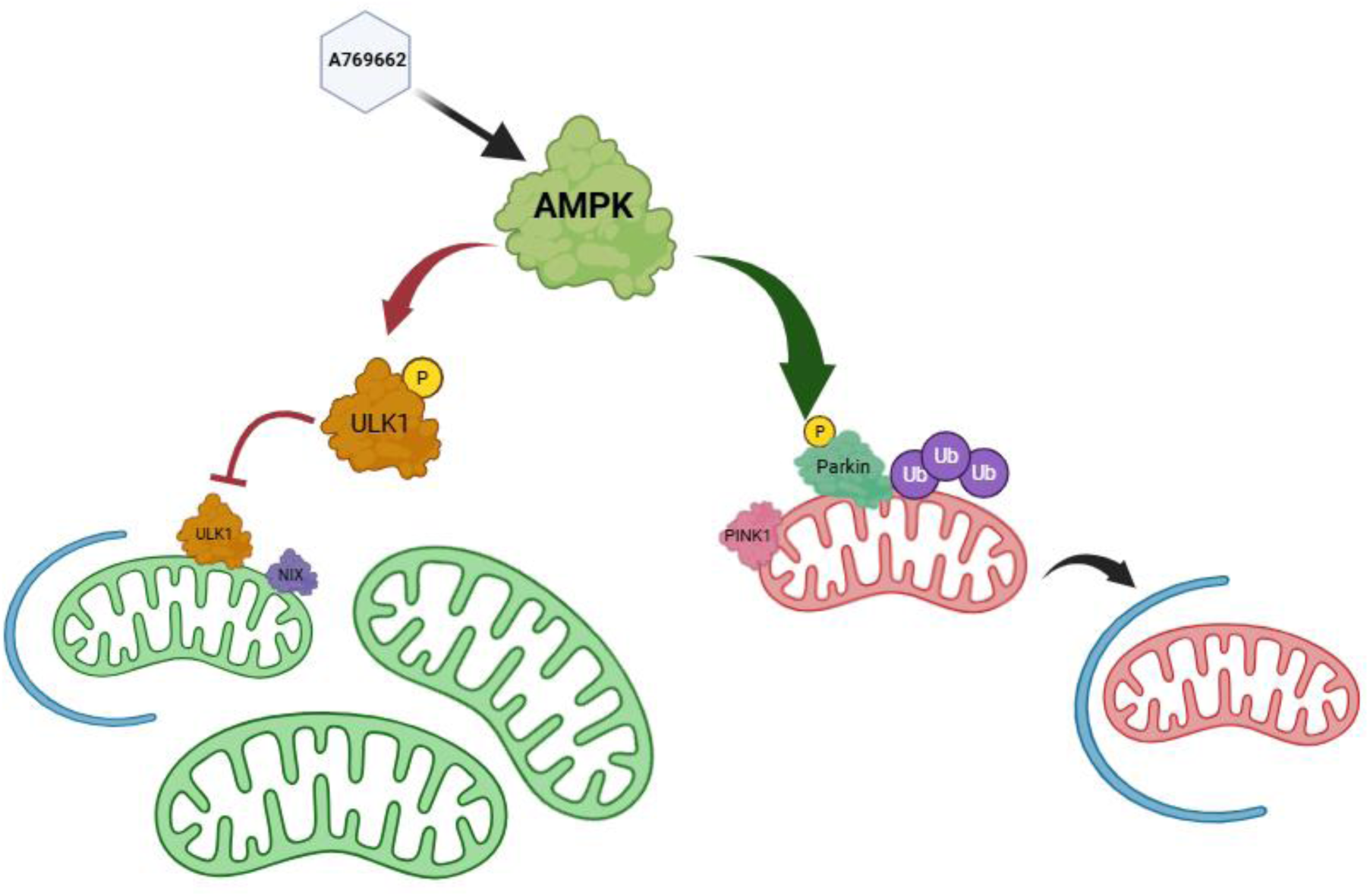
Proposed pathways by which A769662 enhances the proportion of functional mitochondria in LS cells by enhancing overall mitophagy, and in HD cells without altering overall mitophagy levels but altering the balance between mitophagy of functional versus dysfunctional mitochondria.

## Discussion

Mitochondrial dysfunction is a common therapeutic target for many neurological disorders. Very recently the first compound has been approved for use in mitochondrial disease, however this leaves a large unmet need^51^. In this study we used peripheral tissues to characterise mitochondrial health and identify disease relevant phenotypes applicable to phenotypic drug screening. Patient derived fibroblasts are readily accessible and proliferative, making them far more applicable to large drug screening applications that many other human models of disease such as iPSCs. The application of high content imaging provides deep insight in a human disease relevant model, which is scalable to support high throughput screening of thousands of compounds. Using this screening methodology to test tool compounds as proof of concept, we identify a modulator of mitochondrial dysfunction in patient derived fibroblasts from both the primary mitochondrial disorder, LS, and the neurodegenerative disorder, HD.

In LS cells we confirmed mitochondrial phenotypes consisting of reduced respiratory capacity and reduced complex IV activity, as expected based on the published literature^11–13^. However, our data do highlight individual differences between patients and the importance of testing in patient samples. Further to this we were able to detect novel, robust deficits using high throughput imaging in which we identify a significantly reduced proportion of functional mitochondria and reduced mitophagy in LS patient cells. In HD patient derived cells, we identified individual mitochondrial phenotypes in each patient cell line. The phenotypes of LS and HD cells converged on the imbalance between numbers of functional and dysfunctional mitochondria. Our high content imaging of mitochondrial networks enabled us to identify the small molecule A769662, which rescues the imbalance between dysfunctional and functional mitochondria in both LS and HD cells, via differing mechanisms.

Consistent with previous reports, we find reduced mitochondrial oxidative phosphorylation with no corresponding deficit in ATP levels in LS patient cells^13, 52–55^. However, in contrast to Sturm *et al,* we did not find corresponding increases in glycolysis in all LS patient cell lines. Importantly, we identify a lower proportion of mitochondria with a membrane potential in LS patient cells, despite having higher overall MMP, indicating a smaller group of overactive functional mitochondria maintaining a high membrane potential and likely contributing to maintaining ATP levels. We also see a fusion of the functional mitochondrial network in LS cells compared to controls, without a corresponding increase in fusion of the overall network, further suggesting cellular attempts to bolster the functional network. Taking together our respirometry and imaging data, we can build a picture of LS cells with a smaller, more fused functional mitochondrial network than control cells, working harder to maintain ATP levels but still not achieving the same levels of oxidative-phosphorylation as control cells. Notably, we find that LS3 consistently has a less severe mitochondrial phenotype than LS1 and LS2. Higher levels of glycolysis in this line may support mitochondrial function by the enhancing of available substrates. It is also possible that the less severe mitochondrial phenotype could be related to the less severe symptoms reported for this patient, the lack of additional point mutations, or the longer delay between symptom onset and biopsy, potentially having allowed more time for adaptive mechanisms to develop in the patient’s cells.

In galactose culture conditions which drive the need for oxidative phosphorylation, mitochondrial numbers increase in LS cells, yet MMP drops and a lower proportion of the total mitochondria have a membrane potential. This could be driven by fragmentation of damaged mitochondria increasing numbers, or increased biogenesis which inevitably leads to more dysfunctional mitochondria since LS cells with mutations in *SURF1* are less able to assemble proper electron transport chain complexes. Our data suggest LS cells display fragmentation of the dysfunctional parts of the network along with an important fusion of the functional network. Tokuyama et al found that, despite LS cells having more fragmented mitochondria than control cells, inhibiting this fission of the network actually enhances toxicity^53^. This likely indicates that fragmentation of mitochondria in LS cells is related to the need for removal of damaged mitochondria, which complements our finding of reduced percentages of functional mitochondria in all LS cell lines. In accordance with this hypothesis, we report for the first time lower mitophagy flux in LS cells. Over a 6 hour period, especially when mitochondrial damage is stimulated by inhibition of complexes III/V, a smaller proportion of mitochondria are localised to lysosomes in LS cells compared to controls. Moran et al 2014 report larger autophagosomes in fibroblasts from two patients with complex IV deficiency and no corresponding increase in mitochondria at those autophagosomes^56^, also likely indicating a mitophagy deficit. Mitophagy is the subject of much investigation in neurological disorders and many compounds targeting mitophagy have been developed^57^, so a better understanding of the mitophagy deficit in LS could provide novel avenues for drug discovery in LS.

As expected in HD fibroblasts we see low levels of HTT protein and no significant difference in levels between HD and control cells. We find that under basal, glucose containing conditions ATP levels and respiration are consistent between HD and control cells. MMP is significantly reduced in only one patient cell line. In galactose culture conditions, which require the cells to rely more heavily on oxidative phosphorylation, two of three HD cell lines did not show increased ATP levels, however the control cells did.

Our data showing interindividual variability in MMP fits with the fact that reports of basal MMP levels in HD cells are conflicting in the literature^58–61^. Uncoupling conditions highlight MMP, respiratory and complex activity deficits in HD mouse neurons and in other studies of HD patient fibroblasts^61–63^. Jędrak et al found lower ATP levels in HD fibroblasts in both glucose and galactose conditions, however it is unclear if this was a function of the reduced proliferation that they reported in the same cells^64^. Lower ATP levels under oxidative stress conditions are found to be more severe in early-onset compared to late-onset patients^26^. Taken together with these studies, our ATP, MMP and respiratory data together indicate normal respiratory function in HD fibroblasts under basal conditions, with a deficit in the ability to increase oxidative phosphorylation under mitochondrial challenges including lack of glucose and uncoupling conditions.

Our high content imaging studies find an overall phenotype in HD patient derived fibroblasts of enlarged, fused mitochondrial networks, with two of three patient cell lines having a reduced proportion of mitochondria with a membrane potential. Reports in neurons of HD models largely find fragmentation of the mitochondrial network^65–67^, likely indicating cell type specific differences in susceptibility and adaptation to underlying mitochondrial respiratory deficits. Fragmentation of mitochondria is reported in fibroblasts from juvenile HD patients^68,58^, however Vanisova et al (2022) report mitochondrial “swelling” in patient fibroblasts and identify damage to mitochondrial ultrastructure. Higher, sustained co-localisation of mitochondria with lysosomes in HD cells indicates altered mitophagy flux. Since autophagy is well known to be damaged in HD^69^ it is likely that these data represent a stalling of mitochondrial degradation, which would contribute to the higher proportion of dysfunctional mitochondria that we see in HD cells.

The HD patient cells used in this study had comparable CAG repeat lengths (43, 44, 46), thus highlighting individual differences which are not related to the size of repeat expansion. Induced astrocytes from HD patients appear to display more metabolic deficits with higher CAG repeat lengths^70^. Lopes et al found that maximum respiration and spare capacity are in fact increased in pre-manifest patient fibroblasts yet reduced in fibroblasts from patients who have progressed to manifestation of symptoms^27^. Since we find a deficit in HD patient cells to respond to oxidative challenges, it is possible that HD cells initially compensate for underlying mitochondrial deficits but cannot maintain this throughout sustained mitochondrial demand.

### Convergence of phenotypes and effect of AMPK activation

Despite differing underlying causes, and varied forms of mitochondrial dysfunction reported by ourselves and others in LS and HD, we show that cells from both conditions reliably display an imbalance between the number of functional mitochondria and those without a membrane potential. This imbalance could be caused by fusion of the functional network in attempts to maintain activity, or fission or fragmentation of dysfunctional mitochondria, perhaps with concomitant dysfunction in the removal of these damaged mitochondria. Our data suggest fragmentation and corresponding fusion of the functional mitochondrial network in LS and a fusion of the overall network in HD, with mitophagy deficits in cells from both conditions. By applying our high content imaging analysis (through which we identified these mitochondrial phenotypes) to screen a select panel of tool compounds with effects on the mitochondria, we identified the allosteric AMPK activator A769662 as capable of redressing the imbalance between functional and dysfunctional mitochondria present in both LS and HD cells.

AMPK is a major metabolic regulator, sensing AMP/ADP:ATP ratios in order to modulate a multitude of bioenergetic pathways and has attracted much attention in disorders with altered metabolism^47,71^. Altered AMPK levels are reported in fibroblasts from patients with mitochondrial disorders^72^ but have not been reported in LS. AMPK phosphorylates and activates ULK1 which can activate parkin dependent and also independent mitophagy pathways^50^. In response to A769662 treatment, we demonstrated increased ULK1 phosphorylation in control and two of three LS patient cell lines. Relatively low, but highly specific, levels of activation are reported with A769662^73^. AMPK activation has been shown to rescue cell viability and complex activities in cell and mice models of SURF1 deficiency but the mechanisms underlying this rescue are unclear^74,75^. We found that A769662 in LS cells significantly increased the proportion of mitochondria colocalised with lysosomes, indicating an increase in mitophagy consistent with the increase in ULK1 activation. This result might also explain the protective effects of AMPK activation in other SURF1 deficiency models and further highlight mitophagy as a therapeutic target in LS since we have shown that it can be modulated in LS cells and shift the mitochondrial network towards a higher proportion with membrane potential.

AMPK protein levels are reported to be lower in HD mice and induced neurons from HD patients, but not in the fibroblasts from which they were reprogrammed^76,77^, whilst ULK1 activity is reportedly lower in models of HD^78^. A769662 treatment had little effect on AMPK levels but enhanced ULK1 protein or phosphorylation in HD cells. Our findings may suggest that AMPK signalling, although not AMPK protein levels, is altered even in peripheral tissues in HD and these tissues could therefore be used to further study the effects of altered AMPK signalling in HD. Unlike in LS cells, we do not find that A769662 alters overall mitophagy flux in HD cells as measured by co-localisation of mitochondria with lysosomes. There are multiple known pathways of mitophagy, and though their different functions and modulation of them are not fully characterised, recent findings suggest that AMPK can inhibit mitophagy of functional mitochondria via its activation of ULK1 whilst also upregulating parkin dependent mitophagy of dysfunctional mitochondria^79^. It is possible that A769662 treatment and subsequent activation of AMPK and ULK1 induces a shift in mitophagy towards parkin dependent mitophagy, reducing the removal of functional mitochondria without altering overall mitophagy levels.

### Limitations

A key limitation of our study is the small number of patient derived cell lines used. Identifying individual variation is both a strength and a limitation of the use of patient derived cells, but a larger cohort of cell lines would enable better understanding of the individual versus disease common phenotypes. That said, we have identified phenotypes common to all three patient cells which indicates their important in disease. Since HD is known to progress with somatic expansion of the CAG repeat length, it is possible that some dysfunction is masked in HD cells with short CAG repeats, although these repeat lengths are clinically relevant since they are found in patient presenting clinical symptoms. Our tool compound testing only considered five selected compounds and at concentrations which have previously been shown to be effective in other cellular models. In order to fully understand any possible compound effect, a full concentration response curve would need to be undertaken using these and other compounds. In LS cells, we see a relatively modest activation of AMPK with the compound A769662 and future work could investigate the effect of stronger AMPK activators on mitophagy and mitochondrial function in LS cells. In HD cells, although we see a much stronger activation of AMPK compared to their age matched controls, we do not see an activation of mitophagy. Since our study investigates total mitophagy, it is possible that one specific pathway is affected and therefore each pathway would need to be investigated to rule out the effects of A769662 on a single pathway. We have identified an indicator of mitochondrial health, the proportion of mitochondria with intact membrane potential, in both diseases and a small molecule which can rescue deficits in this indicator in fibroblast cells. Limitations to this finding are evidencing that the rescue of functional mitochondria also rescues respiratory dysfunction and that the mechanisms are also relevant in neuronal cells. Further work should seek to understand the effects of AMPK activation on mitochondrial respiration in neuronal models of LS and HD.

### Concluding remarks

To address some of the challenges of drug discovery for complex neurological disorders, we have shown that mitochondrial phenotypes present in LS and HD are identifiable in peripheral tissues, in a high throughput screening format which can be used to identify modifiable parameters. We identify a clear mitophagy deficit for the first time in LS, which could be a valuable therapeutic target. We were able to identify the AMPK activator A769662 as a modulator of both LS and HD relevant mitochondrial features. Together, our work confirms that patient derived fibroblasts recapitulate known disease mechanisms in LS and HD and can be utilised in phenotypic screening to detect small molecule modulators of these phenotypes.

## Contributors

HM led the conceptualisation and experimental design of the study. HM, OB, PJS, LF, SA contributed to securing funding. Experimental work and data analysis was carried out by NH, LE, RH, EM, ES, AT, FE. These authors and HM, GB, AP, OB contributed to data interpretation and presentation. NH and HM contributed to drafting the manuscript to which all authors contributed to reviewing and editing. All authors had full access and approved the final version of the manuscript.

## Acknowledgments

This work was supported by Astellas Pharma Inc. We would like to thank the donors whose cells are stored by the Coriell biorepository.

## Data Sharing Statement

The data reported in this study are available on reasonable request from the corresponding author.

