## Supplementary information for "Identification and small molecule rescue of mitochondrial dysfunction phenotype which converges across Leigh Syndrome and Huntington’s Disease patient fibroblasts"

**Supplementary Table 1:**

*Mitophagy phenotype in HD patient cells and effect of A769662 ^ indicates an increase, v indicates a decrease in HD compared to control (left of table) or in HD cells in response to A769662 treatment (right of table). Data are from live mitophagy assay after 24 hour treatment with DMSO vehicle or 100 µM A769662, in basal conditions and immediately after the addition of 1 µM Oligomycin and 4 µM Antimycin A (Induced).*

**Basal**

| | **HD compared to Control** | | | **A769662 effect** | | |
|---|---|---|---|---|---|---|
| **Parameter** | **HD1** | **HD2** | **HD3** | **HD1** | **HD2** | **HD3** |
| **Mitochondria per cell** | ^ | v | v | v | - | - |
| **Lysosomes per cell** | ^ | v | - | ^ | - | - |
| **% Mitochondria in lysosomes** | - | ^ | - | - | - | - |
| **%Functional mitochondria** | v | v | - | ^ | - | - |
| **MMP** | ^ | - | v | - | - | ^ |

**Induced**

| | **HD compared to Control** | | | **A769662 effect** | | |
|---|---|---|---|---|---|---|
| **Parameter** | **HD1** | **HD2** | **HD3** | **HD1** | **HD2** | **HD3** |
| **Mitochondria per cell** | ^ | - | v | v | v | - |
| **Lysosomes per cell** | ^ | v | v | ^ | - | - |
| **% Mitochondria in lysosomes** | ^ | ^ | - | ^ | - | - |
| **%Functional mitochondria** | ^ | v | v | ^ | - | ^ |
| **MMP** | ^ | - | v | - | - | - |

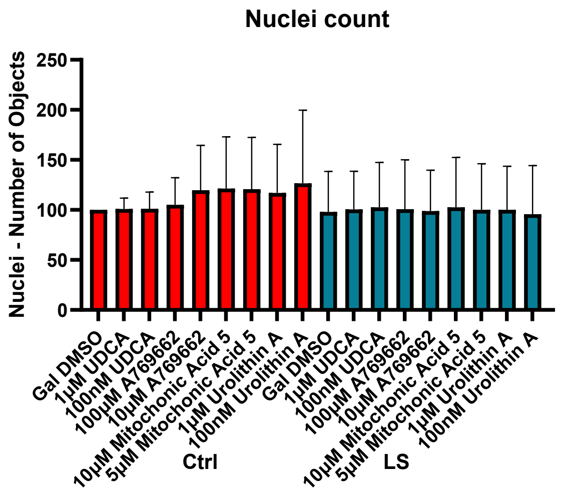


**<u>Supplementary Figure 1:</u>** *Compound effect on cell count in LS patient derived fibroblasts (LS, green) and age matched control cells (Ctrls, red). Effect of 24 hour treatment with 1µM and 100 nM UDCA, 100 µM and 10 µM A769662, 10 and 5 µM MA5 and 1 µM and 100nM UroA on number of individual nuclei. Bars represent the mean of three technical and three biological replicates for each cell line. Error bars represent standard deviation.*

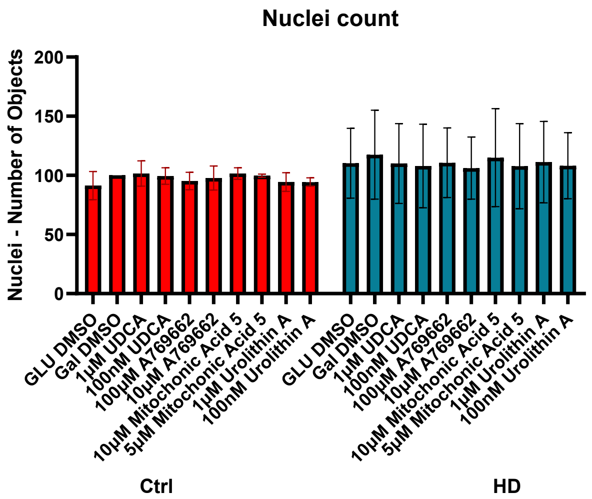


**<u>Supplementary Figure 2:</u>** *Compound effect on cell count in HD patient derived fibroblasts (HD, green) and age matched control cells (Ctrls, red). Effect of 24 hour treatment with 1 µM and 100 nM UDCA, 100 µM and 10 µM A769662, 10 and 5 µM MA5 and 1 µM and 100nM UroA on number of individual nuclei. Bars represent the mean of three technical and three biological replicates for each cell line. Error bars represent standard deviation.*
